# Reprogramming genetic circuits using space

**DOI:** 10.1101/2024.03.20.585869

**Authors:** Lorea Alejaldre, Jesús Miró-Bueno, Angeles Hueso-Gil, Lewis Grozinger, Huseyin Tas, Sina Geißler, Ángel Goñi-Moreno

**Author notes:** These authors contributed equally to this work (Lorea Alejaldre, Jesús Miró-Bueno). Department of Genetics, Harvard Medical School, 77 Avenue Louis Pasteur, Boston, MA 02115.

## Abstract

Genetic circuits confer computing abilities to living cells, performing novel transformations of input stimuli into output responses. Circuit editing often focuses on substituting DNA components, such as RBSs, regulators, or promoters, from part libraries to achieve desired performance. However, this approach is inherently limited by the availability of DNA components. Here, we show that circuit performance can be reprogrammed without altering its DNA sequence by using a library of positions: a set of physical locations within the cell’s volume. Using bacteria as the living chassis, we engineer 219 spatially unique genetic circuits of four different types—three regulatory cascades and a toggle switch—by either inserting the entire circuit in a specific chromosomal position or separating and distributing circuit modules. Their analysis, together with a mathematical model, reveals that spatial positioning can be used not only to optimize circuits but also to switch circuits between modes of operation, giving rise to new functions as circuit complexity increases. We provide foundational insights into leveraging intracellular space for circuit design.

## 1 Introduction

The design and implementation of genetic circuits to process biological information is a central theme in synthetic biology [1, 2, 3, 4]. In these circuits, genetic components receive input signals and output responses are generated based on instructions encoded into gene regulatory networks. This input-algorithm-output pipeline is well aligned with the concept of computation [5], and finds applications in various fields, such as bioproduction [6], pollution control [7] or medical diagnosis [8]. Different host organisms are used for this purpose, often chosen based on the specific application. For example, genetic circuits are now routinely engineered in bacteria [9], yeast [10], plants [11], or mammalian cells [12].

While the selection of DNA parts is crucial for circuit function, and testing libraries of parts is a common practice to fine-tune circuit performance [13, 14], recent years have seen growing interest in the role played by their host context [15, 16, 17]. This refers to the cellular machinery surrounding and interacting with genetic circuits. As a result of these interactions, DNA parts may show different dynamics according to their context, leading to variations in circuit behavior across different cellular hosts [18].

Typically, the consequences of the spatial relationships between genetic circuits and their context are not considered by synthetic biologists. Yet, cells are three-dimensional entities, and even in bacteria (the smallest of them) molecular machineries are not homogeneously distributed throughout the entire cellular volume [19, 20]. Specifically, the chromosome occupies space, and genes are situated at given coordinates. For example, the location of certain genes close to the origin of replication (oriC) or co-localization of co-regulated genes has been proposed to provide a selective advantage for their expression and regulation [21, 22, 23]. It follows that synthetic functions may require a specific address within the volume of the cell—an aspect deserving further attention and that underpins the present work. Ultimately, if evolution has shaped specific spatial configurations for the optimal performance of molecular systems, synthetic biology may find value in incorporating space as a design principle for engineering genetic circuits.

Spatial effects in eukaryotic cells are more extensively described compared to bacteria [24, 25, 26]. Intuitively, one might assume that the presence of physical compartments, higher DNA compaction, and larger volumes in eukaryotic cells makes spatial considerations more significant than in prokaryotes. However, it should not be assumed that space is not a significant factor in bacteria due to their smaller size [27]; indeed, it has been shown that not only are translation and transcription machineries heterogeneously distributed across the cytoplasm [23, 28, 29, 30], but also other types of resources such as enzymes or metabolites are allocated unequally [31]. Moreover, the expression levels of synthetic constructs have been shown to vary based on the specific genomic location in which they are inserted [32], with the common practice being the random integration of the target gene(s) in the genome to select the best performing position [33, 34, 35, 36], therefore using space for optimization. Aside from mean fluorescence levels, circuit performance is also determined by other factors such as gene expression noise which can be conditioned by the distribution of the circuit elements along the sequence [37], the transcription/translation ratio determined by different factors [38], non-genetic causes such as epigenetic modifications [39] or fluctuations in cellular components [40]. All of these factors may be determinant and ultimately influenced by chromosomic position and the availability of resources in that specific point of the three-dimensional cellular space.

The allocation of resources such as ribosomes or polymerases within bacterial cells, crucial for the functioning of synthetic constructs, has strong ties to how the chromosome is organized. For instance, genes relevant to generating these resources tend to be positioned in proximity to the oriC [21, 41]. This specific location confers the advantage of early duplication, resulting in a higher gene dosage throughout the cell lifecycle. Consequently, synthetic constructs located in this area would, in principle, have access to a more robust expression machinery [42, 43]. Other factors, such as DNA supercoiling, upstream transcription or extended protein occupancy domains (EPODs) [44, 45], which are intrinsically space-dependent, can determine how genetic circuits perform depending on the insertion position. Moreover, circuits can be distributed across several locations, the physical separation between them being another spatial aspect with its own dynamics [37].

Here, we argue that space can be effectively harnessed for a new wave of 3D circuits [46]. This becomes even more crucial as circuit complexity increases. Beyond optimizing the expression levels of a single gene, complex circuit behaviors would be intricately affected by spatial modulation. For instance, a basic spatial feature like gene orientation has been shown to play a more significant role in modulating the function of a toggle switch [47] than an expression system [42]. The question that arises is: can it be utilized not only to optimize but also to engineer new biological functions without altering the DNA sequences of synthetic constructs?

In this study, we address this question by using the soil bacterium and synthetic biology chassis *Pseudomonas putida* KT2440 [48, 49, 50, 51]. We randomly inserted and analyzed a variety of gene circuits across different genomic locations to describe the overall positional effects on performance beyond mere expression level modulation. These genetic circuits are flanked by terminators to minimize transcriptional inferences from surrounding genes (Figure S1). Instead of altering DNA sequences, as is commonly done by modifying promoters or ribosome binding sites (RBSs), we manipulated the spatial location of the circuits. In other words, we used location modification as a design tool, comparable to other methods like changing promoter sequences, to modulate circuit function. Following this design strategy, our goal was not to identify the specific factors driving expression changes, but to explore overall positional effects for each genomic location, acknowledging that many factors contribute to variations in circuit behavior. This includes factors such as GC content, the availability of resources like RNAP and ribosomes, transcriptional read-through from upstream genes, chromosomal dynamics, and the physical separation between genes within the same genetic circuit [17, 16, 43, 52]. Our results demonstrate that manipulating the spatial location of circuits can expand their functionality and provide a reference of circuit functionality to aid in the incorporation of spatial design. This expansion ranges from selecting the gene expression noise pattern of individual genes to transforming the functioning of a toggle switch into a sensor switch or a dual-function switch. Based on these findings, we contend that intracellular space can be effectively leveraged to design functions beyond mere sequences, paving the way for a whole-cell design approach in bioengineering.

## 2 Results and discussion

### 2.1 Generation of spatial genetic circuits

To investigate the influence of intracellular space in *Pseudomonas putida* KT2440 on circuit performance, we used a total of 219 genetic variants, each generated by taking one of four genetic constructs and inserting them into the chromosome with a unique spatial configuration. All of our spatial variants are derived from four distinct initial constructs, including three transcriptional cascades and one toggle switch. As a result, our collection of variants has two design features: firstly, the properties of the DNA sequence and genetic components that make up the genetic construct (1D design), and secondly the location and spatial arrangement of these within the chromosome and cell (3D design). Our collection of genetic variants allow us to decouple these two features, so that we can illustrate the potential for spatial variants of identical genetic constructs to impact on circuit behavior (Figure 1A). In other words, we show that while a particular DNA sequence in a specific location and spatial configuration defines a genetic circuit, the circuit can be reprogrammed in various ways, not by changing its DNA sequence, but by using space.

**Figure 1:**
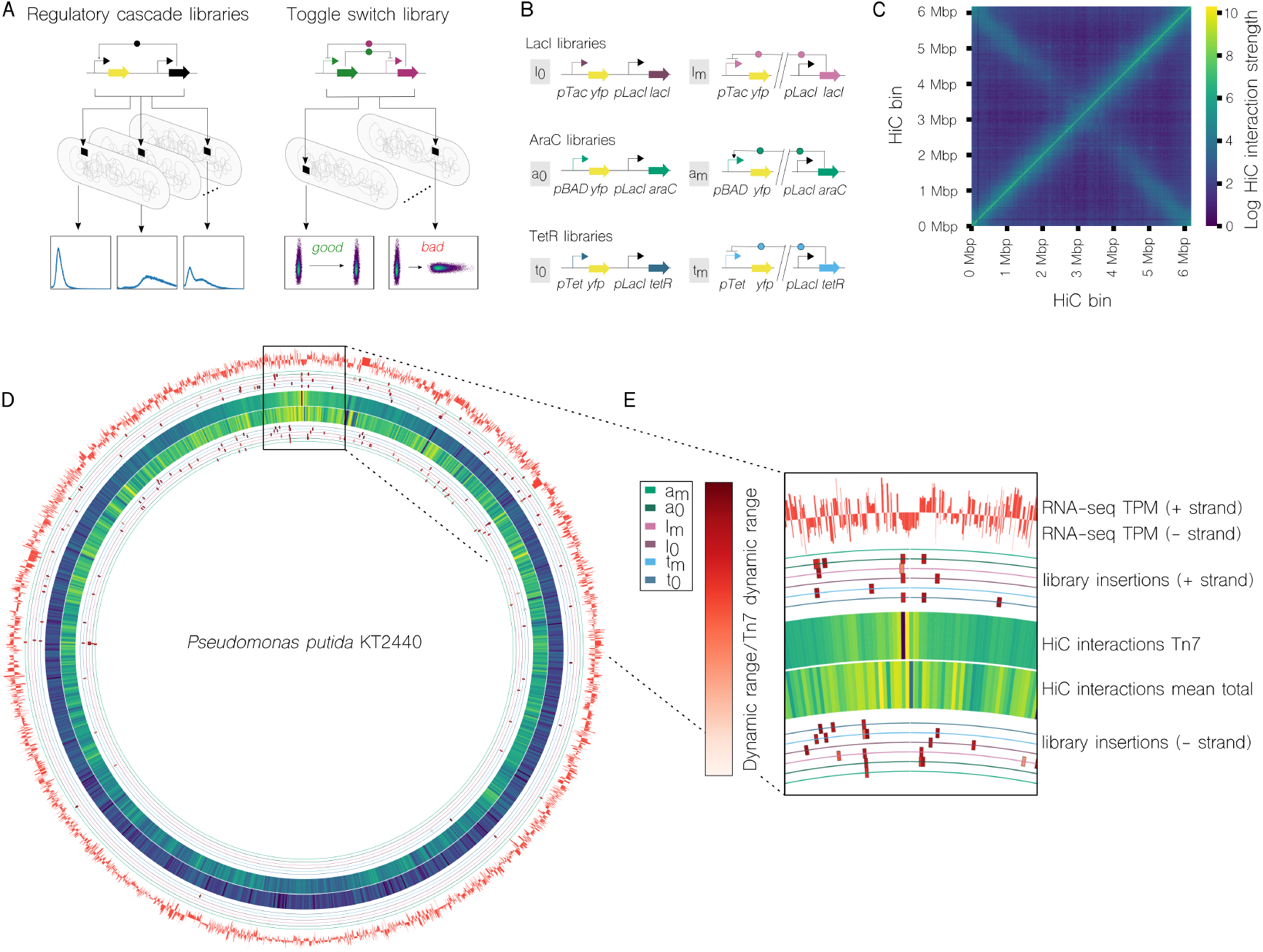
An overview of spatial genetic circuits inserted in a three dimensional *P. putida* chromosome. **A)** We insert two different types of circuit, regulatory cascades and toggle switches, into different chromosomal locations. The phenotype of genetically identical circuits can be reprogrammed by changing where they are inserted. **B)** The three different regulatory cascade circuits used, based on two repressors (LacI and TetR) and one inducer (AraC). For each circuit we tested libraries with all genetic components placed together (l_0_, t_0_ and a_0_), and with the circuit *modularised* and separately distributed across the chromosome (l_*m*_, t_*m*_ and a_*m*_). **C)** HiC contact matrix showing the strength of pairwise interactions between different regions of the *P. putida* KT2440 chromosome (a higher resolution version is included as Figure S2). **D)** HiC interactions alongside RNA-seq analysis of the entire chromosome. Locations where circuits are inserted are marked using a circle for each regulatory cascade library (a higher resolution version is included as Figure S3). **E)** An enlarged section of the full-chromosome of panel D. The six regulatory cascade libraries are shown on colored rings, with boxes marking their insertion position, and the color of the box corresponding to the dynamic range of the circuit compared to that of the same circuit inserted at the Tn7 site.

In the case of our transcriptional cascade constructs, our reprogramming efforts aimed at achieving different output responses that range from expression levels or noise patterns to the dynamic range of circuits (Figure 1A), crucial factors to assess circuit functionality that ultimately influence the compatibility of components within the circuit. Beyond their impact on biological computations, expression noise bears biological significance, such as its potential use in deploying beneficial phenotypes like bet-hedging or cell differentiation strategies [53]. While conventional approaches achieved this by swapping DNA promoters or ribosome binding sites (RBSs), here we have achieved similar outcomes by altering the spatial location.

To further characterize this effect, we constructed six different libraries of transcriptional cascades (Figure 1B). Each library uses one of three different transcription factors, of which two are transcriptional repressors (libraries *l* with LacI and *t* with TetR) and one is a transcriptional activator (*a* with AraC). The libraries can also be divided into two groups based on the spatial configuration of the genetic construct within the chromosome. In the first group, all genetic components of the cascade are inserted together in the same chromosomal position (these libraries are l_0_, t_0_, and a_0_). In the second group, the cascade is split into two distinct modules which are distributed across the chromosome (these libraries are l_*m*_, t_*m*_, and a_*m*_). This separation was conducted such that one module, containing the transcriptional regulator, was consistently positioned at the *attTn7* site to stabilize the level of regulator among variants, while the other module, housing its cognate promoter and the reporter gene, was randomly inserted.

The objective of reprogramming genetic circuits displaying complex phenotypes, such as the bistability exhibited by a toggle switch, is twofold: first, to assess the extent to which spatial positioning is able to modulate the quantitative behavior of the toggle switch, leading to an optimization strategy; and second, we aim to elucidate the emergence of qualitatively new functions, enhancing the reusability of synthetic circuits for different tasks.

We found that all performance variations resulting from distinct spatial positions and arrangements were consistent across replicates and not the consequence of fluctuations or randomness. This reinforces the significance of space as a design layer and indicates fundamental mechanistic differences based on location, essentially forming microenvironments within the intracellular space.

### 2.2 The 3D space of *Pseudomonas putida* KT2440

While *P. putida* KT2440 serves as a widely used chassis in synthetic biology and biotechnology, its intracellular space has only been indirectly studied [29]. To characterize the chromosome of the host cell used in this study, we have generated Hi-C interaction maps and contrasted these with the transcriptional profile based on RNA-seq results, respectively.

The Hi-C contact matrix in Figure 1C depicts the strength of pairwise interactions between chromosomal regions. Aside from featuring a typical diagonal line, indicating strong interactions within a region and its neighbors, focusing on particular rows or columns in this matrix reveals which areas of the chromosome exhibit overall stronger or weaker interactions with the rest of the chromosome —a quality we refer to as the *openness* of a location. The positions of 189 space-specific transcriptional cascade circuits are represented in Figure 1D, with one circular line per library. On each of these lines, inserts are indicated with a square, and the color intensity of each square reflects its performance represented as fluorescence dynamic range normalized to that of the corresponding d=0 circuit inserted in the *attTn7* site for a better comparison between libraries. Concentrically, the outermost circle of Hi-C interactions takes the row in the contact matrix corresponding to the chromosomal region containing the *attTn7* site (of particular interest in this study, as half of the modules in *m* libraries are located there), and represents it as a circular chromosome. Individual sections of this circle are therefore colored according to the interaction strength with the chromosomal region containing the *attTn7* site. The innermost circle of Hi-C interactions in Figure 1D represents openness, with each section colored according to the mean interaction strength of the corresponding chromosomal region with every other region in the chromosome. This figure offers a circular representation of KT2440 chromosome, emphasizing various results of our study. As can be observed in the zoomed-in segment of Figure 1E, interactions with the *attTn7* site are stronger in its vicinity and attenuate as we move farther away in the chromosome. Calculating the openness value for all regions (inner Hi-C circle) reveals substantial variation within the chromosome, even among proximal areas. In essence, certain regions tend to interact more extensively with others, and this characteristic is not uniformly distributed across the chromosome.

To evaluate the influence of transcription activity, we used RNA-seq data previously obtained at the same growth phase as our phenotypic characterization experiments, shown in the outermost circle of Figure 1D [54]. We observed that transcriptional activity and openness profiles do not appear to correlate (Figure S4), suggesting a highly complex relationship between functional layers (i.e., DNA sequence and spatial location). Nevertheless, this chromosomal heterogeneity could offer bioengineers diverse microenvironments to exploit.

Evolution might already be leveraging intracellular space to optimize functions or develop entirely new ones. We compared the co-expressed genomic data of *P. putida* KT2440 against both genomic distance (measured in base pairs) and HiC interaction strength (Figure S5). Results indicate that while most highly co-expressed genes are close together in genomic distance, when considering HiC interaction the opposite trend is observed. Although the discussion of these results is beyond the scope of this work, we believe that the natural utilization of space suggests that a synthetic application of the same is a tool we may want to exploit.

### 2.3 Phenotypic plasticity of transcriptional cascades inserted at different spatial locations

The phenotypic profiles of transcriptional cascades inserted at different genomic positions (all of them can be consulted at the Data availability subsection) were assessed by measuring the reporter’s fluorescence upon induction with either IPTG, aTc or L-Arabinose at saturating concentrations. Flow cytometer measurements revealed variations in circuit performance depending on the spatial location for all the transcriptional cascades (Figures 2A, S6-S11). Whereas variations in expression levels were to be expected based on previous studies [42, 43, 44], here we describe the occurrence of differences in other parameters that impact a synthetic circuit performance such as dynamic range or noise patterns. Considering those terms, our data shows that a better circuit performance is not necessarily linked to higher expression levels (Figures 2C). For example, previous research has proposed that insertion of synthetic circuits at rDNA sites yields higher expression and minimal interference with cellular fitness [35], however this does not guarantee an optimal response to an inducer. In fact, our library variants inserted at rDNA sites show high gene expression but low dynamic range (low dynamic range AraC variants a0B6a, a0F3a, a0G3a and a0H12a in Figure 2C and S12). Therefore, our data shows that the same genetic circuit at a different genomic location can be modulated according to two variables: expression levels and dynamic range. In addition, we note the occurrence of non-standard distributions such as the bimodal distribution (Figure 2A) shown by a few clones from different libraries (more intensely displayed by clones a0C8a and t0H9c, Figures S6 and S10). Among the repertoire of non-canonical behaviors, we also find miss-regulated clones such as tmB6b, t0C8c or t0C4c (Figures S10 and S11) having a permanent induced state even in the absence of aTc. Variations in expression, dynamic range and population distribution for a given variant were found at different genomic regions and chromosomal strands and appear to be highly specific for a given location. For example, the lowest fluorescent variant from the AraC libraries and the highest fluorescent variant from the LacI library have the transcriptional cascade inserted at close positions yet show opposite expression levels (Figure 2A).

**Figure 2:**
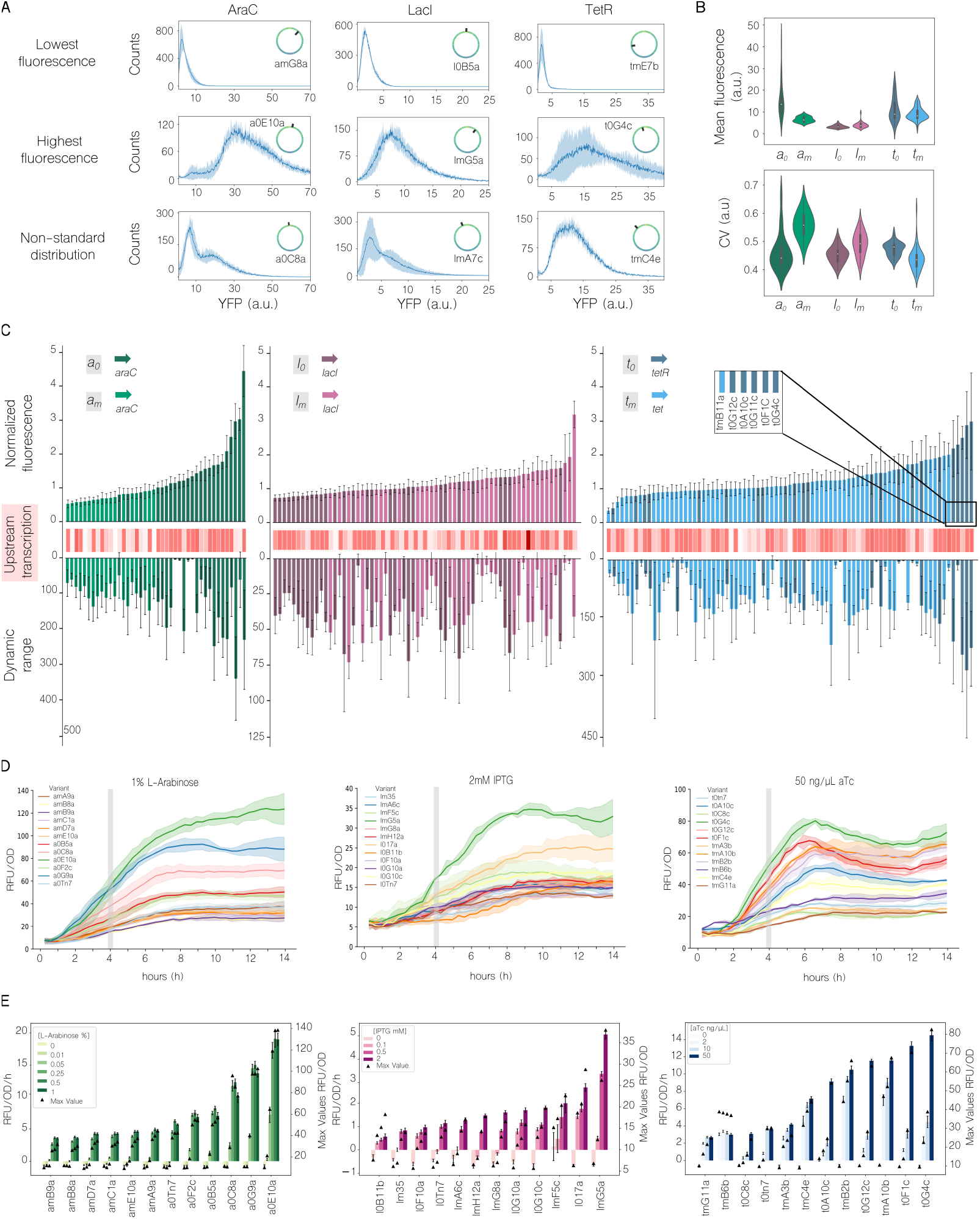
Transposon insertion libraries. **A)** Phenotypic variability in selected individual clones of AraC, LacI and TetR transcriptional cascade insertion libraries. The rest of individual clones behavior can be found at the Figures S6-S11. **B)** Differences between AraC (a_0_ = 23 clones; a_*m*_ = 19 clones), LacI (l_0_ = 31 clones; l_*m*_ = 37 clones) and TetR (t_0_ = 24 clones; t_*m*_ = 57 clones) libraries; mean fluorescence (top) and coefficient of variation (bottom). **C)** Phenotypic characterization of individual clones. Normalized fluorescence to the corresponding transcriptional cascade inserted at *attTn7* for each library (top), transcription of upstream gene for each inserted variant (middle) and dynamic range (bottom). Insertion libraries are ordered as AraC (left), LacI (middle) and TetR (right). (Continued on the following page.) Figure 2: **D)** Fluorescence normalized by growth over time at saturating concentrations of inducer for selected variants of AraC (left), LacI (center) and TetR (right) libraries. **E)** Individual variants performance at different inducer concentrations for AraC (left), LacI (center) and TetR (right) libraries. Maximal fluorescence normalized by growth values appear on secondary y-axis.

By comparing the overall performance of the transcriptional cascade libraries we show differences in the expression levels and noise between them (Figure 2B). In our TetR and LacI transcriptional cascade libraries, we see that separation between the repressor and the cognate promoter showed very little effect on expression levels, compared to when the repressor and cognate promoter were close by (Figure 2B top). However, in our AraC transcriptional cascade library, which is the only transcriptional activator that was integrated in this study, we see a greater effect of distance on expression levels, with the spatially separated a_*m*_ variants showing lower fluorescence on average than the a_0_ constructs. This hints at potential differences between repressors and activators in their effects of space on their dynamics. This is also evident in variants from both AraC libraries that have been inserted in the same spatial location (Figure S13). Another interesting observation is that insertions in position PP 5364 show opposite behaviors for AraC and TetR libraries, lower fluorescence for a_*m*_ library and higher fluorescence for t_*m*_ (Figure S13). The highest expression differences are observed for AraC libraries, whereas the highest variations in dynamic ranges are observed for the TetR library (Figures 2C and S12). While both TetR and LacI are repressors, they show different patterns: LacI shows a lower dynamic range and expression level variation compared to TetR variants. Regarding noise levels we also observe differences between activator and repressor libraries (Figure 2B down). In the activator library, noise levels in a_0_ library are lower compared to a_*m*_. Interestingly, for the two repressors we see two opposing trends. In the LacI library, noise appeared higher in l_*m*_ than in l_0_. Conversely, in the TetR library, the noise in t_*m*_ was lower than in t_0_. Whether these results arise due to the different regulation mechanisms (AraC is an activator; TetR is a repressor) or is particular to these transcription factor/promoter pairs remains to be determined.

The activity of each variant at a given time at saturating concentrations of inducer has revealed that different phenotypic profiles can be obtained for an identical construct at different spatial locations, therefore leading to different operational ranges. Whether this can also be translated to concentrations below saturation or to the behavior over time of the circuits provides an additional layer of complexity in circuit design. 12 variants per transcriptional cascade library, including the *attTn7* control position, were assayed over time and using a range of concentrations below saturation. As seen in Figure 2D, at saturating concentrations of inducer, expression levels reach its maxima between 6-8h post-induction but not at 4 hours, which is the incubation time used in flow cytometry experiments discussed in Figures 2A-C. Considering this, expression levels at 4h do show to be representative of spatial variant performance (Figure 2D). An exception to this is variant a0G9a which shows similar fluorescence over growth at 4h to a0E10a but reaches lower levels of fluorescence. When comparing the performance of variants at concentrations of inducer below saturation (Figure 2E and S14), it is shown that expression levels at saturating concentrations do not necessarily predict expression levels at lower concentrations, obtaining variants with a same genetic circuit that can be modulated differently. In LacI libraries, clones l0G10a and lmG8a show equivalent performance at 2 mM IPTG while at 0.1 mM IPTG lmG8a shows negligible expression of YFP compared to l0G10a. Similarly, variant l0F10a shows higher performance at 0.1 mM IPTG than other variants that reach similar fluorescence values (lm35) or even higher expression at saturating concentrations of IPTG (lmG8a). Several TetR library variants also show this behavior, such as tmG11a which shows higher activity at 10 ng/µL aTc than variants which shows similar expression levels (t0C8c) or higher (t0G12c) at saturating concentrations. Or variants tmA10b and t0G12c which show equivalent performance at 50 ng/µL aTc but t0G12c shows a lower response to concentrations below. Other variants, such as t0F1c, t0G12c and t0A10c display elevated induction at 50 ng/µL aTc but low activity at 10 and 2 ng/µL of aTc (Figure 2E and S15). In addition, the concentration of inducer at which the circuit approaches a saturation status differs in some variants from LacI and TetR libraries but is maintained in AraC library variants which reach saturation at 0.25

Overall, the phenotypic variability observed at different genomic locations suggests a complex scenario where each behavior (considering induced expression and dynamic range) appears to be a consequence of the combination of multiple variables, such as transcription levels of upstream genes, gene dosage due to proximity to the origin of replication or proximity of the transcription factor to its cognate promoter (FigureS16). Although a full mechanistic understanding of the variations in expression levels, dynamic range and noise patterns would offer detailed rules for circuit re-programming using space, our results provide a phenotype-genomic location map reference to use in future strain engineering endeavors which, likewise, greatly aids in 3D design.

### 2.4 Re-programmable genetic switch

To further explore the role of spatial variability in more complex circuits, we next studied a toggle switch combining the two repressors LacI and TetR. Our motivation stems from the variability observed in the transcriptional cascade libraries (Figure 2), which showed marked differences in phenotypes across chromosomal locations. We reasoned that such variability could exert an even more pronounced effect on a more sophisticated network motif like a bistable circuit. To aid this study, we developed both a mathematical model—guided by our previous experimental findings in the transcriptional cascade libraries—and toggle-switch experiments.

#### 2.4.1 Modeling the re-programmability of the genetic switch

We sought to extend our study from the simpler transcriptional cascade libraries to a more complex circuit in which the interplay between multiple repressors could amplify spatial effects. To this end, we developed a mathematical model of a toggle switch composed of LacI and TetR transcription factors. We incorporated two findings from our previous libraries that were particularly relevant for constructing the model. First, we observed that the dynamic range of TetR was consistently higher than that of LacI, suggesting that TetR acts as a stronger repressor in our previous experiments (Figure 2C). In a toggle switch, the system typically features two stable fixed points (representing two expression states A and B) and one unstable fixed point in between [55]. Here, state A corresponds to a high level of TetR and a low level of LacI, whereas state B represents the opposite. When one repressor is stronger, the unstable fixed point shifts closer to the weaker side—here, LacI. Consequently, we chose parameters that place the unstable fixed point nearer to the stable fixed point B. Second, we introduced a noise term *σ*_*r*_ in the model to stochastically perturb each repressor from the state B. Specifically, we used the measured coefficient of variation (CV) from our library data (Figure 2B, bottom) to estimate the range of noise values. (See Methods for model and simulation details.)

Simulations incorporating these findings predict that identical DNA genetic switches can manifest three distinct behaviors, depending on noise strength in different locations (Figure 3). It can function as a toggle switch, where each state remains stable after the inducer is removed (bistability); as a sensor switch, where the system reverts to a single stable state (monostability) upon inducer removal; or as a dual-function switch, where a fraction of the population maintains the induced state and the remainder switches back, giving rise to two coexisting subpopulations. The transition from state A to state B occurs when IPTG is added, resulting in a stable fixed point (green) corresponding to state A (upper row in Figure 3A). Removing IPTG introduces two fixed points: one stable (green) for state B and one unstable (red). If a stable point is very close to an unstable point, noise can push the system towards the unstable point, causing it to jump to the other stable state. The stability of state A is less affected by noise due to the large distance between the stable and unstable fixed points. Conversely, when aTc is added, only the stable fixed point B appears. The reverse transition from state B to state A, shown in the bottom row, indicates that stability of the state B is more affected by noise due to the smaller distance between the points. Exploring the effect of noise strength on the genetic switches (Figure 3B), we found that as noise increases, more cells lose state B and transition to state A, following a sigmoid curve. At low noise, the circuit behaves like a toggle switch, maintaining state B. At intermediate noise, it behaves like a dual-function switch, with some cells in state B and others in state A. At high noise, it functions as a sensor switch, with most cells losing state B.

**Figure 3:**
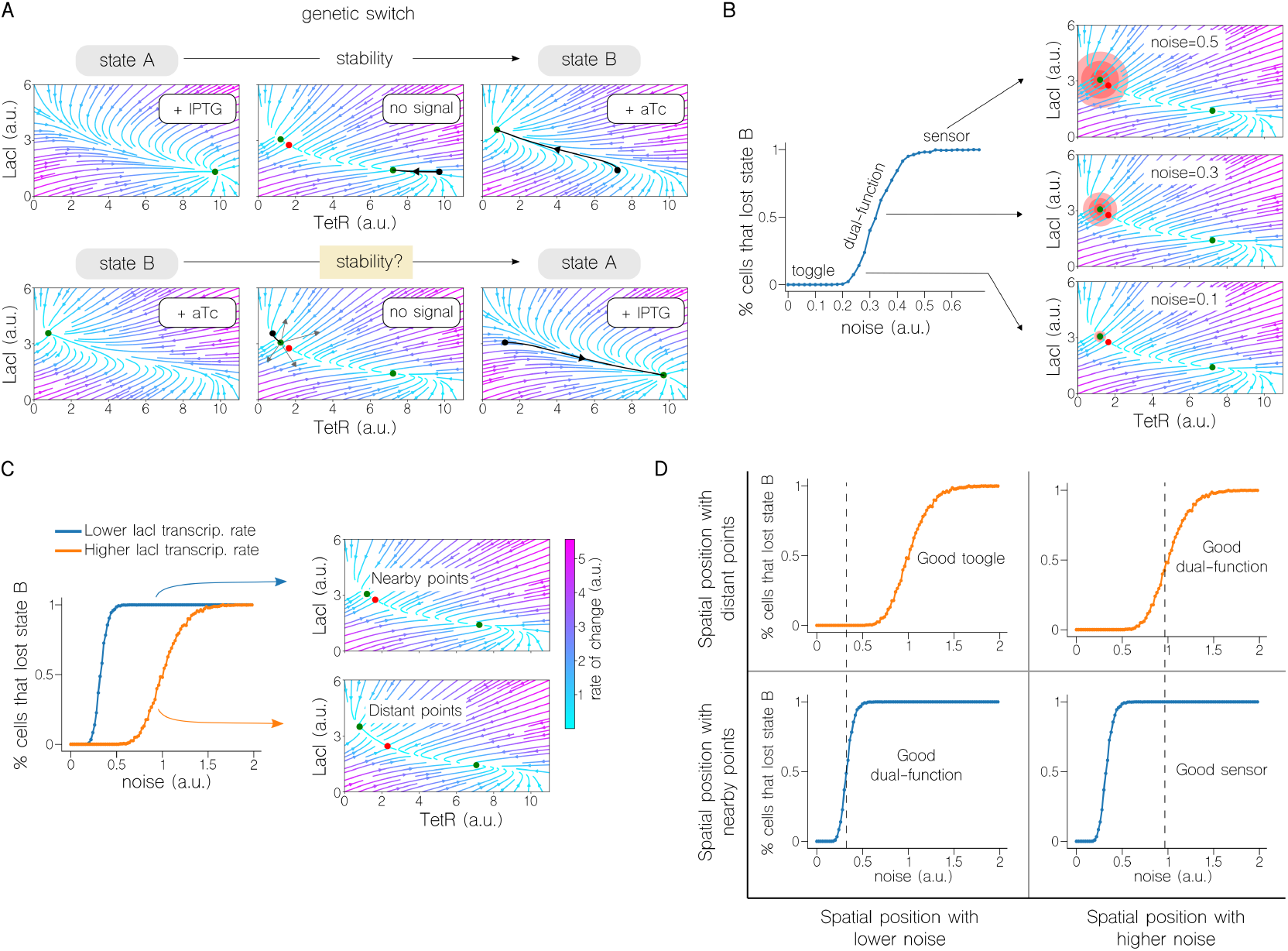
Simulations of the re-programmable genetic switch. **A)** Phase portraits of the re-programmable genetic switch. The top row shows the switch from state A to state B. When IPTG is added, only a stable fixed point (green) appears that corresponds to state A. Removing IPTG introduces two fixed points: one stable (green) for state B, and one unstable (red). Adding aTc results in only the stable fixed point B. The bottom row shows the reverse switch from state B to state A. The stability of state B is more affected by noise after removing aTc due to the small distance between the stable and unstable fixed points, as noise can push the system to the unstable point, causing it to switch to state A. The black line represents the trajectory followed by the fixed point after adding or removing the inducer. The colors of the trajectories represent the rate of change. **B)** Three behaviors depending on noise strength (*σ*_*r*_): at low noise, the circuit behaves like a toggle switch; at intermediate noise, it acts as a dual-function switch; at high noise, it functions as a sensor switch. The percentage of cells losing state B as a function of noise strength follows a sigmoid curve. Three circles, with radii of *σ*_*r*_, 2*σ*_*r*_, and 3*σ*_*r*_, are included for visual reference of noise strength for each *σ*_*r*_ value. **C)** Two different sigmoidal curves depending on spatial positions with different basal transcription rates of lacI. Higher basal transcription rates result in stable and unstable fixed points being close, yielding a short region for toggle and dual-function behaviors and a long region for sensor behavior. Lower basal transcription rates result in the fixed points being farther apart, showing a longer region for toggle and dual-function behaviors and a shorter region for sensor behavior.Figure 3: The function of the genetic switch depends on noise and the distance between the fixed points in each spatial position. Lower noise and shorter distance (or higher noise and greater distance) improve dual-function switch performance. Higher noise and shorter distance enhance sensor switch performance. Lower noise and greater distance favor toggle switch performance. The orange line in the upper graphs shows simulations with more separated points, while the blue line in the bottom graphs shows simulations with closer points. The dashed line is a reference for comparing the orange and blue curves with the same noise strength. (See Methods for model and simulation details).

Finally, incorporating variables that shift the repressors’ strength balance—i.e., those that change the position of the unstable fixed point, such as the basal transcription rate—underscores how different chromosomal positions can alter whether the circuit remains bistable, becomes monostable, or bifurcates into two subpopulations. We observed that noise affected the sigmoidal curves differently depending on the spatial position of the genetic switches (Figure 3C). For instance, positions with higher basal transcription rates of the gene lacI resulted in stable and unstable fixed points being close together, yielding a short region for the toggle and dual-function switch behaviors and a long region for the sensor behavior. Conversely, lower basal transcription rates led to stable and unstable fixed points being farther apart, resulting in a longer region for the toggle and dual-function switch behaviors and a shorter region for the sensor behavior. From these results we can conclude that the function of the genetic switch depends on both the noise and the distance between the stable and unstable fixed points in each spatial position (Figure 3D). There are four possible combinations: lower noise and shorter distance (or higher noise and greater distance) result in better performance as a dual-function switch; higher noise and shorter distance enhance performance as a sensor switch; lower noise and greater distance improve performance as a toggle switch.

Taken together, these simulations provided essential guidelines for interpreting the subsequent experimental results, in which relocating the same toggle switch circuit in different spatial positions of *P. putida* indeed yielded all three predicted behaviors. These findings also guided our subsequent experimental analysis, providing the rationale for identifying toggle-like, sensor-like, and dual-function phenotypes (see Methods for details on the scoring functions).

#### 2.4.2 Phenotypic validation of a reprogrammable toggle switch

Out of the 219 engineered circuits, 30 were spatial variants of a genetic toggle switch (Figure 4A and Table S1). Importantly, the DNA sequence of the circuit remained unchanged across all variants, with the variations arising solely from differences in spatial positioning and the intrinsic constraints of specific locations. Given the constant DNA sequence, the toggle switches may be expected to display two states in all cases: either expressing green or red fluorescence. These states imply the activation of the circuit towards the *msfGFP* (state A) or *mKate2* (state B) genes, respectively, while remaining inactive in the other direction. To induce these states, we used the same two signals: IPTG for turning on state A and aTc for turning on state B. The definition and identifiability of the states is therefore the first basic fingerprint of this circuit. Notably, the stability of each state after removing the inducers represents the second fundamental feature of this bistable switch.

Regarding the definition of states, we characterized circuits by assessing populations of each toggle switch variant after inducing with IPTG or aTc. This approach accounted for expression noise and intrinsic variability in both green and red fluorescent values. The states of the circuits were then evaluated based on population activity, aiming for clear distinctions between states without overlaps. In other words, when in state A, the activity of the population should be distinguishable from when the circuit was in state B. Figure 4B illustrates a well-defined separation, demonstrating that the area covered by cells expressing msfGFP (state A) does not overlap with the area covered by the same cells when expressing mKate2 (state B). The same figure also highlights instances of poorly-defined states, where a significant portion of the population expresses both genes simultaneously at different levels. This leads to circuits with suboptimal or even useless performance. Consequently, a toggle switch with poorly-defined states may possess a well-designed DNA sequence inadequately located within the chromosome. In such cases, altering the DNA parts is unnecessary; instead, relocating the circuit to a different location is the key solution-as demonstrated below through the classification of the 30 switch variants based on their performance.

**Figure 4:**
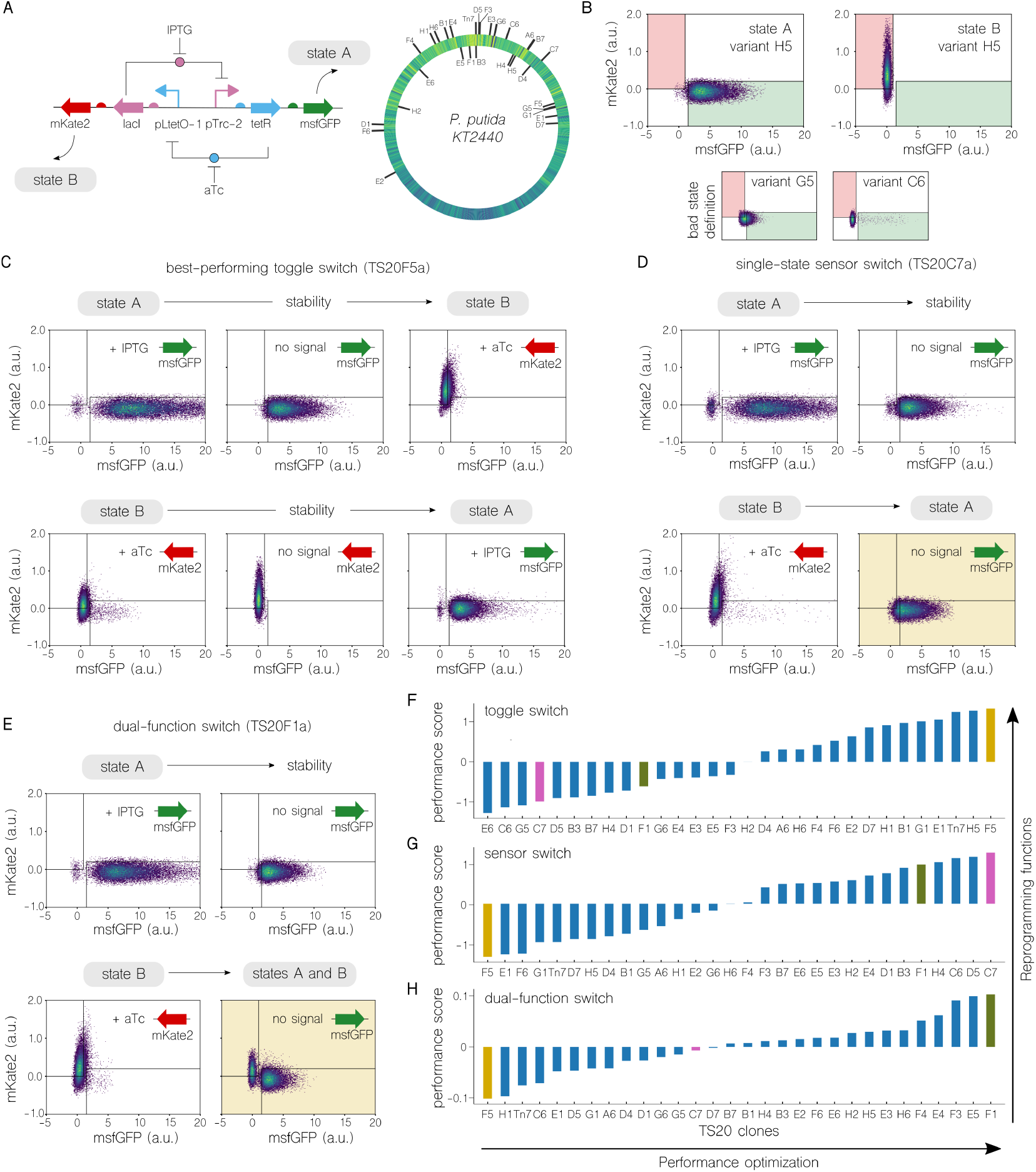
Re-programmable genetic switch. **A)** Schematic of the toggle switch (left). The genetic circuit comprising *tetR* and *lacI* genes, coding for repressor proteins TetR and LacI. These proteins suppress each other’s expression, creating a toogle switch. Output is via mKate2 and msfGFP fluorescent proteins, regulated by TetR and LacI, respectively. The circuit transitions between states A and B with IPTG or aTc inducers. IPTG inactivates LacI, enabling *tetR* and msfGFP expression (state A), while aTc inactivates TetR, allowing *lacI* and mKate2 expression (state B). The right side of the panel illustrates the location of the 30 spatial variants of the genetic toggle switch within the Hi-C ring map of *P. putida* KT2440.Figure 4:**B)** Density plots illustrating states A and B of the toggle switch for variant TS20H5a. The top-left plot demonstrates state A, with the cell population predominantly located within the green square. The top-right plot shows state B, where most cells fall within the red square. The two bottom plots present instances of suboptimal switching states for variants TS20G5a and TS20C6a, where the cell populations predominantly fall outside the designated squares, highlighting an undesirable or ‘bad’ state configuration. **C)** Performance of variant TS20F5a demonstrated through the transition dynamics between states A and B. The top row depicts the switch from state A to B, beginning with 4 hours of IPTG induction to achieve state A, followed by maintaining state A stability after IPTG removal for 20 hours, and concluding with a switch to state B after 4 hours of aTc induction. The bottom row reverses this process from state B to A. **D)** Performance of the variant TS20C7a as a sensor switch. The system maintains stability post-IPTG induction (top row), but loses stability and reverts to state A after aTc removal (bottom row), highlighting its differential response to inducers. **E)** Performance of the variant TS20F1a as a dual-function switch. The genetic circuits maintain the state after the induction of IPTG (top row). After aTc removal, the cell population divides into two subpopulations: one part maintains state B, while the other part moves to state A (bottom row). **F), G)**, and **H)** Performance scores of the 30 variants as toggle, sensor and dual-function switches (see Methods for the score calculation). The best toggle switch variant, TS20F5a, performs poorly as a sensor switch and dual-function switch, while the top sensor switch, TS20C7a, and the top dual-function switch, TS20F1a, rank low as toggle switches.

On the other hand, reprogramming the stability of states can lead to the emergence of new functions. Within the library of 30 switches, some exhibit stability in one state but not in the other, while still preserving the ideal state definition. Stability refers to the ability to maintain a state once reached, regardless of whether the inducing signal that triggered the activation of that state is still present. A bistable switch, like the variant TS20F5a (Figure 4C), exemplifies this performance. After inducing with IPTG to activate state A, the signal can be removed, and the circuit retains the state permanently until the other inducing signal, aTc, turns off state A and triggers the activation of state B. Similarly, state B is also stable. In contrast, variant TS20C7a (Figure 4D) lacks stability in state B. Consequently, once in state B, the circuit autonomously transitions to reach state A as soon as aTc is removed, rendering it unstable. A potential mechanistic explanation, consistent with our model’s prediction, is that an imbalance in the repression strengths of TetR and LacI, which can vary depending on the chromosomal location, shifts the unstable fixed point closer to the LacI-dominant regime. This asymmetry, together with noise—whose magnitude can also vary with location—destabilizes state B after aTc removal, allowing the system to transition back toward TetR dominance. Such a monostable switch could be useful in scenarios where a default state is needed, and the other state will only be triggered temporarily while its sensing signal is present—function what we refer to as the sensor switch. Another distinct behavior is shown by the variant TS20F1a (Figure 4E), in which the population is divided into two states, A and B, after aTc removal. A potential mechanistic explanation for this behavior, again consistent with our model, is that the imbalance in repressor strengths at this particular chromosomal location situates the unstable fixed point in such a way that, in combination with the noise at this location, cells can be pushed toward either state A or B. Consequently, once aTc is removed, not all cells converge on a single state; instead, part of the population transitions to state A while another fraction remains in state B, yielding two coexisting subpopulations. This performance, which we call the dual-function switch, could be interesting in situations where one signal makes the system perform one task, while the other signal makes the system perform two tasks simultaneously.

As stated before, a well-designed circuit can still underperform if placed in a suboptimal chromosomal location; thus, we next classified the 30 switch variants based on their toggle switch performance (Figure 4F), revealing significant variability within the library, with approximately 50% passing our quality test (see Methods). Furthermore, our results suggest that poorly performing variants should not be discarded outright. For example, variants TS20C7 and TS20F1a, which show poor toggle switch performance, function very well as a sensor switch (Figure 4G) and as a dual-function switch (Figure 4H), respectively. Although no definitive rules emerged from the genetic switch library (Figures S17), the distinct behaviors across positions highlight the versatility of space as a design parameter, suggesting that further specialized exploration could uncover underlying constraints.

## 3 Conclusions

Living systems have evolved intricate mechanisms that we have yet to fully understand. Nevertheless, synthetic biology has successfully integrated many of these mechanisms into its design toolkit. However, the spatial flow of information within living cells, meticulously shaped by evolution, is crucial for gene regulation and has yet to be fully incorporated. This research formalizes space as a critical design feature within that toolkit. Just as spatial considerations are fundamental in electronic circuit design, they are also vital in genetic circuit engineering but often overlooked. This study underscores the importance of spatial positioning and provides information to utilize it in bioengineering efforts.

By engineering 219 unique spatial configurations of four types of genetic circuits, including regulatory cascades and toggle switches, this study shows how circuit behavior can change due to spatial positioning, stressing the utility of this type of positional maps. In the regulatory cascades, not only circuit performance as dynamic range can be tuned using space, but we also managed to precisely control gene expression patterns, ranging from full expression to bimodal activity and beyond. Building upon insights from these simpler cascade libraries, we developed a mathematical model to explore a more intricate circuit: the toggle switch, in which the interplay between multiple repressors can amplify spatial effects. Selecting an appropriate position for a complex circuit may lead to the exploration of new functionalities: we experimentally transformed toggle switches into sensor switches or dual-function switches without modifying their DNA sequence. While the experiments and model presented here cannot fully resolve all the mechanistic details underlying the 219 spatial configurations, they do offer valuable insights. For instance, in the toggle switch, we show how variations in repressor strengths and noise levels may shape the different observed phenotypes. Although we show here how to build predefined functions using space even without full mechanistic knowledge, further studies on intracellular spatial dynamics will deepen our understanding of, and control over, these spatially driven phenomena.

This work paves the way to reprogramming complex features by using spatial positioning, such as the stability of circuit states. Location modification can be used just like any other part of the circuit: modifying the promoter sequence or the spatial arrangement are two comparable options (though distinct) to modulate circuit function. The engineering of biological functions without altering DNA sequences significantly expands the design space for genetic circuits. These circuits could be optimized based on a target function or entirely reprogrammed to achieve new operational modes.

## 4 Material and Methods

### Strains, media and general culture conditions

LB media was used to culture both *E. coli* and *P. putida* strains except when M9 minimal media supplemented with 0.2% (v/w) citrate was required: during certain steps of the conjugation protocol according to previous references ([56]; [57]) and for the cultivation of library clones for flow cytometry. Antibiotics were used at the following concentrations: kanamycin (Km) 50 µg/mL, chloramphenicol (Cm) 50 µg/mL, gentamicin (Gm) 10 µg/mL, and ampicillin (Ap) 150 µg/mL for *E. coli*, while 500 µg/mL for *P. putida. E. coli* and *P. putida* strains were cultured at 37ºC and 30ºC respectively. For every specific library, inducers were added at the following concentrations: Isopropyl *β*-D-1-thiogalactopyranoside (IPTG) 2 mM, anhydrotetracyclin (aTc) 50 ng/µL, and L-arabinose (L-ara) 1%. The list of strains used in this work is shown in Table 1. *P. putida* KT2440 was the receiver strain for every integration event. A set of *E. coli* strains coding for Pir protein were used as parental strains in order to keep suicide vectors (pBAMD1.2, pBLAM1.2 and pTn7-M derivatives built in this study; Table 2). *E. coli* HB101 bearing pRK600 was used as the helper strain for all integrations, while *E. coli* DH5*αλ*pir transformed with pTNS2 plasmid (carrying a transposase) assisted mini-Tn7 transpositions.

**Table 1:**
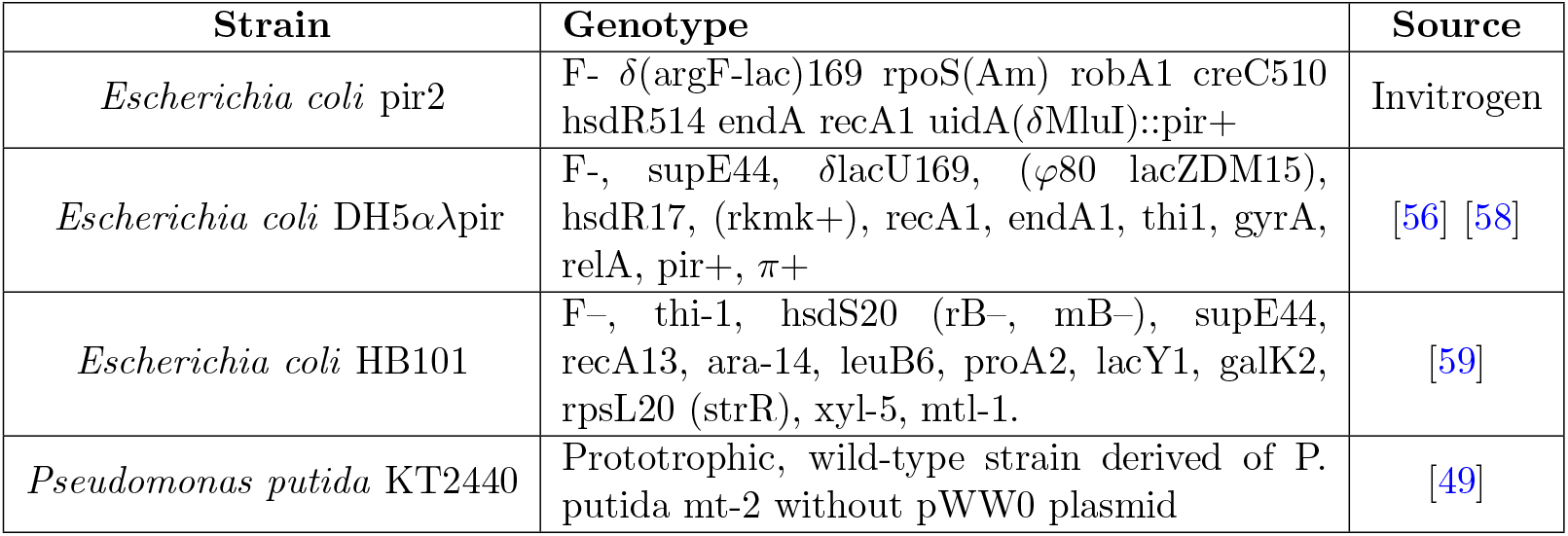
List of strains used in this study.

**Table 2:**
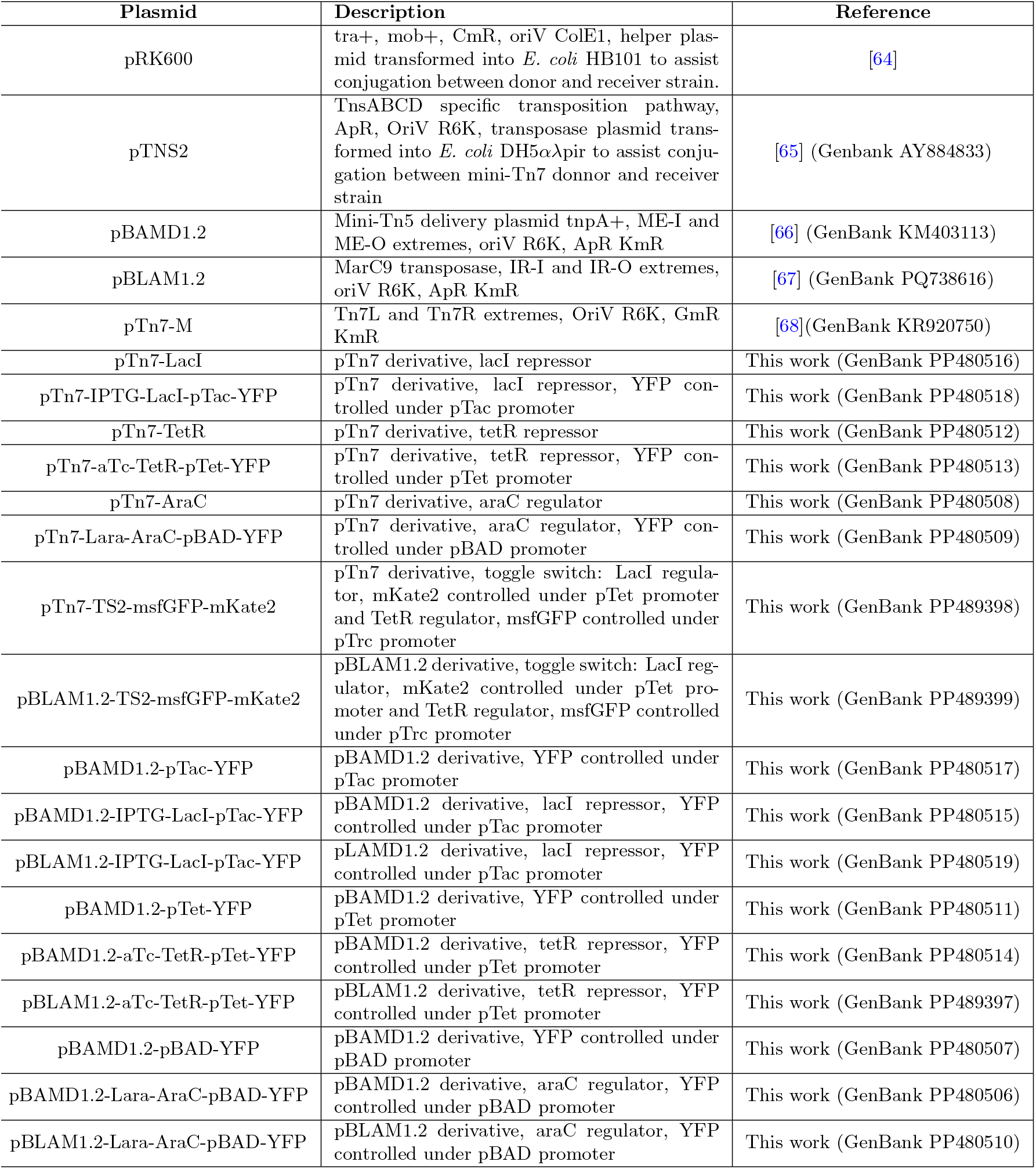
List of plasmids used in this work.

### DNA assemblies

Plasmids used and built in this study are listed in Table 2. Genetic circuit sequences aTc-TetR-pTet, IPTG-LacI-pTac, 3MB-XylS-pM, Lara-AraC-pBAD were ordered from Doulix (Rome, Italy) and Toggle Switch Class 2 (TS2) fragments TS2-TetR-msfGFP and TS2-LacI-mKate2 from Twist Bioscience (California, USA). TS2 fragments were assembled to generate the toggle switch circuit via a touchdown SOE-PCR protocol from [60]. DNA was cloned using either classical restriction and ligation ([61]; restriction enzymes, T4 DNA ligase and Quick Ligation kit from New England Biolabs, Ipswich, Massachusetts, USA) or isothermal assembly ([62]; Gibson Assembly MasterMix, New England Biolabs, Ipswich, Massachusetts, USA). Plasmid pBLAM1-2-TS2-msfGFP-mKate2 was subjected whole-plasmid site-directed mutagenesis[63] to eliminate MmeI site in the TetR gene. DNA oligos (Table S2) were ordered to IDT (Coralville, Iowa, USA) and polymerase enzymes used for DNA amplification were Phusion (Thermo Fisher Scientific, Waltham, Massachusetts, USA) or Q5 (New England Biolabs, Ipswich, Massachusetts, USA). The construction method for each plasmid built in this work can be consulted in Table S3. Minipreps were performed using Monarch Plasmid Miniprep kit (New England Biolabs, Ipswich, Massachusetts, USA) and PCR products were purified with Monarch PCR and DNA Cleanup kit (New England Biolabs, Ipswich, Massachusetts, USA).

### Construction, selection, and genotyping of transposon insertion libraries

Once pTn7-M, pBAMD1.2 and pBLAM1.2 derivative suicide plasmids were built, they were delivered into *P. putida* KT2440 chromosome following a conjugation protocol previously described for transpositions into gram-negative bacteria [56] [57]). Conjugation time was reduced to 5h in order to minimize replication of variants or spurious events, such as multiple genomic insertions. Conjugation outcome was processed in order to separate variants, discard variants with spurious integrations and genotype the transposon location in accordance with a previously described automated high-throughput screening method [67] that uses LAP protocols [69] and is also available through protocols.io[70]. Selected and genotyped libraries were stored at -80ºC for further phenotypic characterization. The genomic location of the inserted genetic circuits separated by library can be consulted in the Data availability subsection.

### Flow cytometry

96-well flat-bottomed plates were filled with 100 µL of LB per well plus the proper antibiotic and inoculated with selected variants previously stored at -80ºC. This preinoculum was placed at 30ºC with shaking for an O/N incubation. The following day, 1 µL of the preinoculum was used to inoculate a new plate filled with 100 µL of filtered M9 media supplemented with 0.2% (v/w) citrate plus the proper antibiotic and the corresponding inducer at saturating concentrations (2 mM IPTG, 1% L-Arabinose or 100 ng/µL aTc). This plate was incubated at 30ºC with 500 rpm shaking for 4 hours. After incubation, 150 µLof M9 citrate were added in order to dilute cultures and balance the events per second required for an adequate flow cytometry measurement. For the Toggle switch variants, after 4h induction with either 100 ng/µL aTc, 2 mM IPTG or not inducer, 20 µL were diluted with 100 µL M9 citrate and measured in the flow cytometer; the remaining 80 µL of culture were diluted in 800 µL of M9 citrate, centrifuged at 2204 × g and resuspended in 100 µL of M9 citrate to wash chemical inducers. Toggle switch cultures were grown at 30ºC with shaking for 20h without inducers, 10 µL of cultures were diluted with 100 µL of M9 citrate and measured in the flow cytometer and 1 µL used to inoculate 100 µL of M9 citrate with the opposite inducer (IPTG for variants previously induced with aTc or aTc for those induced with IPTG) or no inducer. After 4h of growth at 30ºC with shaking, cultures were measured in the flow cytometer. Cultures were passed through a MACSQuant VYB Flow Cytometer (Miltenyi Biotec, Germany) at a maximum speed of 20,000 events per second up to 100,000 singlet events. All variants were measured at least in triplicate. To clean the cytometry raw data, we applied several gates to each sample. Outliers were removed across all channels using the z-score parameter. Next, gating was applied to the FSC-A and SSC-A density plots, and doublets were eliminated using gating on the SSC-H versus SSC-A density plot. Samples that displayed abnormal SSC-H versus SSC-A density plots, resulting from running through the cytometer at rates exceeding maximum speed, were excluded from the analysis. After data cleaning, 20,000 events from each sample were randomly selected for the final statistical analysis. Bar-graph values represented correspond to the fluorescence mean normalized to that of the d=0 circuit at the *attTn7* site. Dynamic range values were obtained by dividing mean fluorescence upon induction by mean fluorescence in the absence of induction.

### Individual variant fluorescence and growth assay in 96-well plate format

12 genomic variants (6 X_0_ library clones including the *attTn7* site variant, and 6 X_*m*_ library clones) per transcriptional cascade library were individually assayed over time and between several concentrations using a Varioskan LUX multimode microplate reader. Selected variants were inoculated from glycerol stocks into 5 mL of Luria-Bertani media. After overnight growth at 30ºC, preinoculums were set to an OD600nm =10 and 2 µL were inoculated in 200 µL of M9 citrate with a range of inducer concentrations. AraC variants were assayed at 0, 0.01, 0.05, 0.25, 0.5 and 1% L-Arabinose; LacI variants were assayed at 0, 0.1, 0.5 and 2 mM IPTG and TetR variants at 0, 2, 10 and 50 ng/µL. All variants were inoculated twice in a 96-well plate to perform technical duplicates. The same layout was repeated in separate experiments to obtain at least three biological replicates. Growth was measured as turbidimetric scattering at 600 nm and YFP fluorescence was measured with wavelengths of excitation *λ*_*Ex*_ = 485 nm and emission *λ*_*Em*_ = 535 nm. Growth and fluorescence were measured with a time lapse of 20 minutes during a minimum of 14h at 28ºC. Transcriptional cascade variant performance was measured as the slope in the linear range of the increase in relative fluorescence units (RFU) normalized by growth over time. The linear range for all variants was established between 140 min (2,3 h) and 400 min (6,7 h). Slopes were determined using an ordinary least squares linear regression model.

### RNAseq data analysis

RNASeq raw data of replicates of *P*.*putida* KT2440 in exponential phase grown in M9-citrate liquid media were obtained from [54]. To calculate transcription expression levels, data was processed using Geneious prime 2023.2.1 (Biomatters Ltd, New Zealand). Reads were aligned to the reference genome of *P*.*putida* KT2440 (Accession number NC 002947.3) using the built-in Geneious RNA mapper. Transcripts per million (TPM) for reads mapped against the annotated reference genome were calculated using the built-in function “Calculate expression levels”.

### HiC data acquisition and analysis

*P*.*putida* KT2440 and *P*.*putida* KT-GFP strains were grown overnight from a glycerol stock at 30ºC with shaking. These precultures were inoculated into 5 mL M9-citrate at OD600nm of 0.1 and grown at 30ºC for 4h. Cultures were then centrifuged, fixed with formaldehyde following Phase Genomics (Seattle, Washington, USA) guidelines and sent for HiC processing (DNA extraction, digestion with Sau3AI and MluCI restriction enzymes and sequencing). HiC reads QC analysis was performed by Phase Genomics. Each culture was sent in duplicate. Analysis of HiC reads in .fastq format was performed using the TADbit software pipeline, built as a Singularity image from a modified specification that is made publicly available on github. The pipeline used is as follows:

1. Index the reference genome using gem3-mapper.
2. Map reads to the reference genome using gem3-mapper.
3. Parse and filter reads using the default filters employed by the TADbit filter tool.
4. Assign filtered reads into bins of 5000 bp
5. Normalise bins using the “Vanilla” normalization offered by TADbit (a single iteration ICE normalization [71]).
6. Convert bin list to matrix format.

More details of the analysis can be obtained by inspecting the script we used to run the pipeline, which is publicly available on github and contains the specific arguments given to TADbit for each step. This github repository also contains the scripts and data files used to generate the sub panels of Figure 1. Quality control plots are provided in the supplementary information (Figures S18-S19).

### Model of the genetic switch

The mathematical model of the genetic switch consists of two dimensionless differential equations:

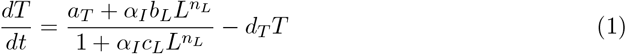

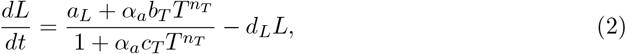

where *T* and *L* are the amounts of TetR and LacI, respectively. The parameters *a*_*L*_ and *a*_*T*_ primarily account for the basal transcription rates of LacI and TetR. The parameters *b*_*L*_ and *b*_*T*_ primarily account for the reduction in the transcription rates of LacI and TetR due to repression. The parameters *c*_*L*_ and *c*_*T*_ account for the cooperativity and oligomerization rates of LacI and TetR. The parameters *d*_*L*_ and *d*_*T*_ represent the degradation rates of LacI and TetR. The exponents *n*_*L*_ and *n*_*T*_ represent the tetramerization of LacI and the dimerization of TetR, respectively. The parameters *α*_*I*_ and *α*_*a*_ account for the inhibition effects of the inducers IPTG and aTC on LacI and TetR, respectively. Specifically, *α*_*I*_ modifies the values of the parameters *b*_*L*_ and *c*_*L*_, while *α*_*a*_ modifies the values of the parameters *b*_*T*_ and *c*_*T*_. The model presented above is based on biochemical reactions that describe gene expression and regulation [55, 72].

We incorporated two observations from the libraries into the mathematical model. First, our library data suggest that TetR is stronger than LacI in this particular chromosomal context (Figure 2C). If one repressor is stronger than the other, the unstable fixed point will shift toward the “weaker” side. Accordingly, we chose parameters that place the unstable fixed point closer to the stable fixed point representing the state B. For the simulations shown in Figure 3, the parameters used were *a*_*T*_ = 5, *a*_*L*_ = 1.8, *b*_*L*_ = 0.01, *b*_*T*_ = 0.12, *c*_*L*_ = 0.1, *c*_*T*_ = 0.2, *d*_*L*_ = 0.5, *d*_*T*_ = 0.5, *n*_*L*_ = 4, *n*_*T*_ = 2, *α*_*I*_ = 1 and *α*_*a*_ = 1. The orange curves in Figure 3 were calculated with *a*_*L*_ = 1.9. When IPTG is added, *α*_*I*_ is set to 0.1, and when aTc is added, *α*_*a*_ is set to 0.1.

Second, we used the coefficient of variation (CV) measurements from the libraries (Figure 2B, bottom) to simulate noise. Specifically, we perturbed the system from the fixed point B, denoted as (*T*_*B*_, *Y*_*B*_), using random values generated from normal distributions. The noise strength is defined as *σ*_*r*_. To account for noise in both axes, we express it as 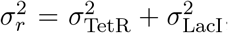, where 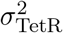 and 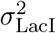 are the standard deviations of the normal distributions 𝒩(0, *σ*_TetR_) and 𝒩(0, *σ*_LacI_). Because the observed CV values for l_0_ and t_0_ libraries were comparable, we set *σ*_LacI_ = *σ*_TetR_ = *σ*, so that 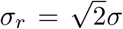. In the model, we perturb the system from its fixed point by *σ*_*r*_, which we set to the order of magnitude of the CV in our data. Therefore, if Δ*T* ∼ 𝒩(0, *σ*) and Δ*L* ∼ 𝒩(0, *σ*), the position of the system was perturbed to (*T*_*B*_ + Δ*T, L*_*B*_ + Δ*L*). We then checked whether the system remained at fixed point B or moved to fixed point A. For calculating each point in the blue and orange curves in Figure 3, we repeated this process for 1000 cells.

### Score calculation

For the calculation of the toggle switch performance, we used the following equations:

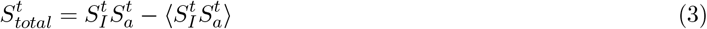

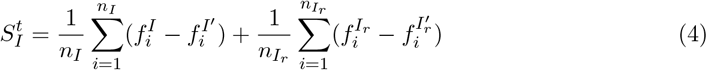

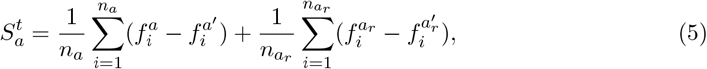

where 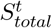 is the total performance score, and 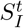 and 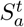 are the scores for IPTG and aTC inducers, respectively. 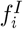 and 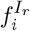 represent the fraction of events in the green square after IPTG induction and removal, while 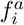 and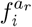 denote the fraction in the red square after aTc induction and removal, respectively. 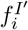 and 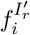 represent the fraction of events in the red square after IPTG induction and removal, while 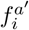 and 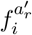 denote the fraction in the green square after aTc induction and removal, respectively. *n*_*I*_ and *n*_*a*_ are the numbers of replicates for IPTG and aTc inductions, while *n*_*Ir*_ and *n*_*ar*_ are the numbers of replicates after IPTG and aTc removal, respectively.

For the calculation of the sensor switch performance, we used the following equations:

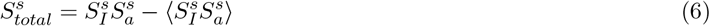

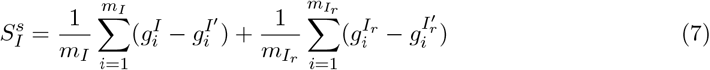

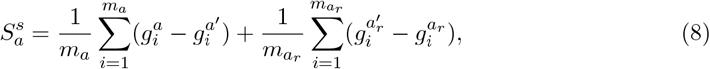

where 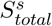 is the total performance score, and 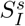 and 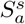 are the scores for IPTG and aTC inducers, respectively. 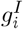 and 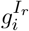 represent the fraction of events in the green square after IPTG induction and removal, while 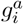 and 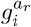 denote the fraction in the red square after aTc induction and removal, respectively. 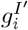 and 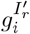 represent the fraction of events in the red square after IPTG induction and removal, while 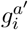 and 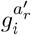 denote the fraction in the green square after aTc induction and removal, respectively. *m*_*I*_ and *m*_*a*_ are the numbers of replicates for IPTG and aTc inductions, while *m*_*Ir*_ and *m*_*ar*_ are the numbers of replicates after IPTG and aTc removal, respectively.

For the calculation of the dual-function switch performance, we used the following equations:

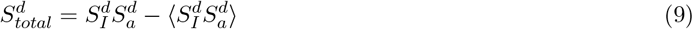

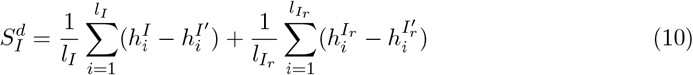

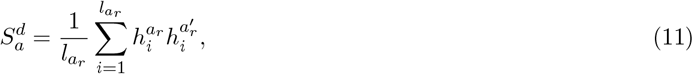

where 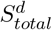 is the total performance score, and 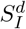 and 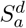 are the scores for IPTG and aTC inducers, respectively. 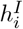 and 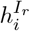 represent the fraction of events in the green square after IPTG induction and removal, while 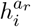 denotes the fraction in the red square after aTc removal. 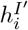 and 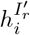 represent the fraction of events in the red square after IPTG induction and removal, while 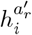 denotes the fraction in the green square after aTc removal. *l*_*I*_ is the number of replicates for IPTG induction, while *l*_*Ir*_ and *l*_*ar*_ are the numbers of replicates after IPTG and aTc removal, respectively.

All experiments involving the toggle library were conducted in quadruplicate. Only samples that passed the flow cytometry gating criteria were considered for the score calculation. Density plots illustrating the performance of all clones are available in the Data availability subsection.

## Data availability

The flow cytometry data used for analysis in this study, the information regarding the positions and statistics of all the variants in the regulatory cascade libraries, and the density plots of all the switch variants showing their performance are available here.

## Acknowledgments

This work was supported by the the ECCO (ERC-2021-COG-101044360) Contract of the EU, grants BioSinT-CM (Y2020/TCS-6555) and CONTEXT (2019-T1/BIO-14053)of the Comunidad de Madrid, and grants MULTI-SYSBIO (PID2020-117205GA-I00), BIOELECTRIC (CNS2022-135951), and MULTISYNBIO (PID2023-152470NB-I00) funded by MICIU/AEI/ 10.13039/501100011033.

## Notes

### Competing Interest Statement

The authors have declared no competing interest.

### Summary of Updates

Added: mathematical modelling, predictions, library curation, time-based experiments, supplementary info with more details on insulation and individual performance.

## References

[1] Y. Benenson, “Biomolecular computing systems: principles, progress and potential,” Nature Reviews Genetics, vol. 13, no. 7, pp. 455–468, 2012.

[2] E. Andrianantoandro, S. Basu, D. K. Karig, and R. Weiss, “Synthetic biology: new engineering rules for an emerging discipline,” Molecular systems biology, vol. 2, no. 1, pp. 2006–0028, 2006.

[3] M. Amos and A. Goñi-Moreno, “Cellular computing and synthetic biology,” Computational matter, pp. 93–110, 2018.

[4] J. Miró and A. Rodríguez-Patón, “Biomolecular computing devices in synthetic biology,” International Journal of Nanotechnology and Molecular Computation (IJNMC), vol. 2, no. 2, pp. 47–64, 2010.

[5] L. Grozinger, M. Amos, T. E. Gorochowski, P. Carbonell, D. A. Oyarzún, R. Stoof, H. Fellermann, P. Zuliani, H. Tas, and A. Goñi-Moreno, “Pathways to cellular supremacy in biocomputing,” Nature communications, vol. 10, no. 1, p. 5250, 2019.

[6] V. Chubukov, A. Mukhopadhyay, C. J. Petzold, J. D. Keasling, and H. G. Martín, “Synthetic and systems biology for microbial production of commodity chemicals,” NPJ systems biology and applications, vol. 2, no. 1, pp. 1–11, 2016.

[7] V. De Lorenzo, K. L. Prather, G.-Q. Chen, E. O’Day, C. von Kameke, D. A. Oyarzún, L. Hosta-Rigau, H. Alsafar, C. Cao, W. Ji, et al., “The power of synthetic biology for bioproduction, remediation and pollution control: the un’s sustainable development goals will inevitably require the application of molecular biology and biotechnology on a global scale,” EMBO reports, vol. 19, no. 4, p. e45658, 2018.

[8] S. Slomovic, K. Pardee, and J. J. Collins, “Synthetic biology devices for in vitro and in vivo diagnostics,” Proceedings of the National Academy of Sciences, vol. 112, no. 47, pp. 14429–14435, 2015.

[9] A. A. Nielsen, B. S. Der, J. Shin, P. Vaidyanathan, V. Paralanov, E. A. Strychalski, D. Ross, D. Densmore, and C. A. Voigt, “Genetic circuit design automation,” Science, vol. 352, no. 6281, p. aac7341, 2016.

[10] M. W. Gander, J. D. Vrana, W. E. Voje, J. M. Carothers, and E. Klavins, “Digital logic circuits in yeast with crispr-dcas9 nor gates,” Nature communications, vol. 8, no. 1, p. 15459, 2017.

[11] T. K. Kassaw, A. J. Donayre-Torres, M. S. Antunes, K. J. Morey, and J. I. Medford, “Engineering synthetic regulatory circuits in plants,” Plant Science, vol. 273, pp. 13–22, 2018.

[12] M. Mansouri and M. Fussenegger, “Therapeutic cell engineering: designing programmable synthetic genetic circuits in mammalian cells,” Protein & Cell, vol. 13, no. 7, pp. 476–489, 2022.

[13] B. A. Blount, T. Weenink, S. Vasylechko, and T. Ellis, “Rational diversification of a promoter providing fine-tuned expression and orthogonal regulation for synthetic biology,” PloS one, vol. 7, no. 3, p. e33279, 2012.

[14] R. G. Egbert and E. Klavins, “Fine-tuning gene networks using simple sequence repeats,” Proceedings of the National Academy of Sciences, vol. 109, no. 42, pp. 16817–16822, 2012.

[15] A. Boo, T. Ellis, and G.-B. Stan, “Host-aware synthetic biology,” Current Opinion in Systems Biology, vol. 14, pp. 66–72, 2019.

[16] C. Moschner, C. Wedd, and S. Bakshi, “The context matrix: Navigating biological complexity for advanced biodesign,” Frontiers in Bioengineering and Biotechnology, vol. 10, 2022.

[17] S. A. Scholz, C. D. Lindeboom, and P. L. Freddolino, “Genetic context effects can override canonical cis regulatory elements in Escherichia coli,” Nucleic Acids Research, vol. 50, pp. 10360–10375, Oct. 2022.

[18] H. Tas, L. Grozinger, R. Stoof, V. de Lorenzo, and Goñi-Moreno, “Contextual dependencies expand the re-usability of genetic inverters,” Nature Communications, vol. 12, p. 355, Jan. 2021. Number: 1 Publisher: Nature Publishing Group.

[19] I. V. Surovtsev and C. Jacobs-Wagner, “Subcellular organization: a critical feature of bacterial cell replication,” Cell, vol. 172, no. 6, pp. 1271–1293, 2018.

[20] J. Kim, A. Goñi-Moreno, and V. de Lorenzo, “Subcellular architecture of the xyl gene expression flow of the tol catabolic plasmid of pseudomonas putida mt-2,” mBio, vol. 12, no. 1, pp. 10–1128, 2021.

[21] A. Soler-Bistúe, M. Timmermans, and D. Mazel, “The Proximity of Ribosomal Protein Genes to oriC Enhances Vibrio cholerae Fitness in the Absence of Multifork Replication,” mBio, vol. 8, pp. e.00097–17, Feb. 2017.

[22] E. Garmendia, G. Brandis, and D. Hughes, “Transcriptional Regulation Buffers Gene Dosage Effects on a Highly Expressed Operon in Salmonella,” mBio, vol. 9, pp. e01446–18, Sept. 2018.

[23] J. Fan, H. El Sayyed, O. J. Pambos, M. Stracy, J. Kyropoulos, and A. N. Kapanidis, “RNA polymerase redistribution supports growth in E. coli strains with a minimal number of rRNA operons,” Nucleic Acids Research, vol. 51, pp. 8085–8101, Aug. 2023.

[24] M. M. Babu, S. C. Janga, I. de Santiago, and A. Pombo, “Eukaryotic gene regulation in three dimensions and its impact on genome evolution,” Current opinion in genetics & development, vol. 18, no. 6, pp. 571–582, 2008.

[25] S. Leidescher, J. Ribisel, S. Ullrich, Y. Feodorova, E. Hildebrand, A. Galitsyna, S. Bultmann, S. Link, K. Thanisch, C. Mulholland, et al., “Spatial organization of transcribed eukaryotic genes,” Nature cell biology, vol. 24, no. 3, pp. 327–339, 2022.

[26] D. B. Brückner, H. Chen, L. Barinov, B. Zoller, and T. Gregor, “Stochastic motion and transcriptional dynamics of pairs of distal DNA loci on a compacted chromosome,” Science, vol. 380, pp. 1357–1362, 2023.

[27] M. Campos and C. Jacobs-Wagner, “Cellular organization of the transfer of genetic information,” Current Opinion in Microbiology, vol. 16, pp. 171–176, Apr. 2013.

[28] M. Irastortza-Olaziregi and O. Amster-Choder, “Coupled Transcription-Translation in Prokaryotes: An Old Couple With New Surprises,” Frontiers in Microbiology, vol. 11, p. 624830, Jan. 2021.

[29] J. Kim, A. Goñi-Moreno, B. Calles, and V. de Lorenzo, “Spatial organization of the gene expression hardware in Pseudomonas putida,” Environmental Microbiology, vol. 21, no. 5, pp. 1645–1658, 2019. Section: 1645.

[30] S. A. Scholz, R. Diao, M. B. Wolfe, E. M. Fivenson, X. N. Lin, and P. L. Freddolino, “High-Resolution Mapping of the Escherichia coli Chromosome Reveals Positions of High and Low Transcription,” Cell Systems, vol. 8, pp. 212–225.e9, Mar. 2019.

[31] K. M. Dahlstrom and G. A. O’Toole, “A Symphony of Cyclases: Specificity in Diguanylate Cyclase Signaling,” Annu Rev Microbiol, vol. 71, pp. 179–195, Sept. 2017.

[32] S. Köbbing, T. Lechtenberg, B. Wynands, L. M. Blank, and N. Wierckx, “Reliable Genomic Integration Sites in Pseudomonas putida Identified by Two-Dimensional Transcriptome Analysis,” ACS Synthetic Biology, vol. 13, pp. 2060–2072, July 2024. Publisher: American Chemical Society.

[33] T. E. Saleski, M. T. Chung, D. N. Carruthers, A. Khasbaatar, K. Kurabayashi, and X. N. Lin, “Optimized gene expression from bacterial chromosome by high-throughput integration and screening,” Science Advances, vol. 7, p. eabe1767, Feb. 2021.

[34] A. Hueso-Gil, B. Calles, and V. de Lorenzo, “In Vivo Sampling of Intracellular Heterogeneity of Pseudomonas putida Enables Multiobjective Optimization of Genetic Devices,” ACS Synthetic Biology, May 2023.

[35] A. Domröse, J. Hage-Hülsmann, S. Thies, R. Weihmann, L. Kruse, M. Otto, N. Wierckx, K.-E. Jaeger, T. Drepper, and A. Loeschcke, “Pseudomonas putida rDNA is a favored site for the expression of biosynthetic genes,” Scientific Reports, vol. 9, p. 7028, Dec. 2019.

[36] J. E. Chaves, R. Wilton, Y. Gao, N. M. Munoz, M. C. Burnet, Z. Schmitz, J. Rowan, L. H. Burdick, J. Elmore, A. Guss, D. Close, J. K. Magnuson, K. E. Burnum-Johnson, and J. K. Michener, “Evaluation of chromosomal insertion loci in the Pseudomonas putida KT2440 genome for predictable biosystems design,” Metabolic Engineering Communications, vol. 11, p. e00139, Dec. 2020.

[37] Á. Goñi-Moreno, I. Benedetti, J. Kim, and V. de Lorenzo, “Deconvolution of gene expression noise into spatial dynamics of transcription factor–promoter interplay,” ACS synthetic biology, vol. 6, no. 7, pp. 1359–1369, 2017.

[38] E. M. Ozbudak, M. Thattai, I. Kurtser, A. D. Grossman, and A. van Oudenaarden, “Regulation of noise in the expression of a single gene,” Nat Genet, vol. 31, pp. 69–73, May 2002. Publisher: Nature Publishing Group.

[39] A. Sanchez, S. Choubey, and J. Kondev, “Regulation of noise in gene expression,” Annu Rev Biophys, vol. 42, pp. 469–491, 2013.

[40] D. J. Kiviet, P. Nghe, N. Walker, S. Boulineau, V. Sunderlikova, and S. J. Tans, “Stochasticity of metabolism and growth at the single-cell level,” Nature, vol. 514, pp. 376–379, Oct. 2014. Publisher: Nature Publishing Group.

[41] M. Wolański, R. Donczew, A. Zawilak-Pawlik, and J. Zakrzewska-Czerwińska, “oriC-encoded instructions for the initiation of bacterial chromosome replication,” Frontiers in Microbiology, vol. 5, 2015.

[42] D. H. S. Block, R. Hussein, L. W. Liang, and H. N. Lim, “Regulatory consequences of gene translocation in bacteria,” Nucleic Acids Research, vol. 40, no. 18, pp. 8979–8992, 2012. Section: 8979.

[43] T. E. Kuhlman and E. C. Cox, “Gene location and dna density determine transcription factor distributions in Escherichia coli,” Molecular Systems Biology, vol. 8, no. 1, p. 610, 2012.

[44] J. A. Bryant, L. E. Sellars, S. J. W. Busby, and D. J. Lee, “Chromosome position effects on gene expression in Escherichia coli K-12,” Nucleic Acids Research, vol. 42, pp. 11383–11392, 09 2014.

[45] V. Gerganova, M. Berger, E. Zaldastanishvili, P. Sobetzko, C. Lafon, M. Mourez, A. Travers, and G. Muskhelishvili, “Chromosomal position shift of a regulatory gene alters the bacterial phenotype,” Nucleic Acids Research, vol. 43, pp. 8215–8226, Sept. 2015.

[46] Z. Li, H. Yang, Y. Wang, S.-H. Chou, and J. He, “The spatial position effect: synthetic biology enters the era of 3D genomics,” Trends in Biotechnology, vol. 40, pp. 539–548, 2022.

[47] E. Yeung, A. J. Dy, K. B. Martin, A. H. Ng, D. Del Vecchio, J. L. Beck, J. J. Collins, and R. M. Murray, “Biophysical constraints arising from compositional context in synthetic gene networks,” Cell Systems, vol. 5, no. 1, pp. 11–24.e12, 2017.

[48] E. Martínez-García and V. de Lorenzo, “Pseudomonas putida as a synthetic biology chassis and a metabolic engineering platform,” Current Opinion in Biotechnology, vol. 85, p. 103025, 2024.

[49] K. E. Nelson, C. Weinel, I. T. Paulsen, R. J. Dodson, H. Hilbert, V. a. P. Martins dos Santos, D. E. Fouts, S. R. Gill, M. Pop, M. Holmes, L. Brinkac, M. Beanan, R. T. DeBoy, S. Daugherty, J. Kolonay, R. Madupu, W. Nelson, O. White, J. Peterson, H. Khouri, I. Hance, P. C. Lee, E. Holtzapple, D. Scanlan, K. Tran, A. Moazzez, T. Utterback, M. Rizzo, K. Lee, D. Kosack, D. Moestl, H. Wedler, J. Lauber, D. Stjepandic, J. Hoheisel, M. Straetz, S. Heim, C. Kiewitz, J. Eisen, K. N. Timmis, A. Düsterhöft, B. Tümmler, and C. M. Fraser, “Complete genome sequence and comparative analysis of the metabolically versatile Pseudomonas putida KT2440,” Environmental Microbiology, vol. 4, no. 12, pp. 799–808, 2002.

[50] H. Tas, L. Grozinger, A. Goñi-Moreno, and V. de Lorenzo, “Automated design and implementation of a nor gate in pseudomonas putida,” Synthetic Biology, vol. 6, no. 1, p. ysab024, 2021.

[51] H. Tas, Á. Goñi-Moreno, and V. d. Lorenzo, “A standardized inverter package borne by broad host range plasmids for genetic circuit design in gram-negative bacteria,” ACS synthetic biology, vol. 10, no. 1, pp. 213–217, 2020.

[52] R. T. Dame, F.-Z. M. Rashid, and D. C. Grainger, “Chromosome organization in bacteria: mechanistic insights into genome structure and function,” Nature Reviews Genetics, vol. 21, pp. 227–242, Apr. 2020.

[53] J.-C. García-Betancur, A. Goñi-Moreno, T. Horger, M. Schott, M. Sharan, J. Eikmeier, B. Wohlmuth, A. Zernecke, K. Ohlsen, C. Kuttler, et al., “Cell differentiation defines acute and chronic infection cell types in staphylococcus aureus,” Elife, vol. 6, p. e28023, 2017.

[54] Hueso-Gil, B. Calles, G. A. O’Toole, and V. De Lorenzo, “Gross transcriptomic analysis of Pseudomonas putida for diagnosing environmental shifts,” Microbial Biotechnology, vol. 13, pp. 263–273, Jan. 2020.

[55] T. S. Gardner, C. R. Cantor, and J. J. Collins, “Construction of a genetic toggle switch in Escherichia coli,” Nature, vol. 403, no. 6767, pp. 339–342, 2000.

[56] E. Martínez-García, B. Calles, M. Arévalo-Rodríguez, and V. de Lorenzo, “pBAM1: an all-synthetic genetic tool for analysis and construction of complex bacterial phenotypes,” BMC Microbiology, vol. 11, p. 38, Feb. 2011.

[57] E. Martínez-García and V. de Lorenzo, “Engineering multiple genomic deletions in Gram-negative bacteria: analysis of the multi-resistant antibiotic profile of Pseudomonas putida KT2440,” Environmental Microbiology, vol. 13, no. 10, pp. 2702–2716, 2011.

[58] D. Hanahan, “Studies on transformation of Escherichia coli with plasmids,” Journal of Molecular Biology, vol. 166, pp. 557–580, June 1983.

[59] J. J. Schmidt, “DNA cloning: A practical approach,” Biochemical Education, vol. 14, no. 2, pp. 190–245, 1986.

[60] R. S. Hilgarth and T. M. Lanigan, “Optimization of overlap extension PCR for efficient transgene construction,” MethodsX, vol. 7, p. 100759, Jan. 2020.

[61] J. Sambrook, E. F. Fritsch, and T. Maniatis, Molecular Cloning: A Laboratory Manual. Cold Spring Harbor Laboratory, 1989.

[62] D. G. Gibson, L. Young, R.-Y. Chuang, J. C. Venter, C. A. Hutchison, and H. O. Smith, “Enzymatic assembly of DNA molecules up to several hundred kilobases,” Nat Methods, vol. 6, pp. 343–345, May 2009.

[63] M. Laible and K. Boonrod, “Homemade site directed mutagenesis of whole plasmids,” J Vis Exp, May 2009.

[64] B. Kessler, V. de Lorenzo, and K. N. Timmis, “A general system to integrate lacZ fusions into the chromosomes of gram-negative eubacteria: regulation of the Pm promoter of the TOL plasmid studied with all controlling elements in monocopy,” Mol Gen Genet, vol. 233, pp. 293–301, May 1992.

[65] K.-H. Choi, J. B. Gaynor, K. G. White, C. Lopez, C. M. Bosio, R. R. Karkhoff-Schweizer, and H. P. Schweizer, “A Tn7-based broad-range bacterial cloning and expression system,” Nat Methods, vol. 2, pp. 443–448, June 2005.

[66] E. Martínez-García, T. Aparicio, V. de Lorenzo, and P. I. Nikel, “New Transposon Tools Tailored for Metabolic Engineering of Gram-Negative Microbial Cell Factories,” Frontiers in Bioengineering and Biotechnology, vol. 2, 2014.

[67] L. Alejaldre, A.-M. Anhel, and Goñi-Moreno, “pBLAM1-x: standardized transposon tools for high-throughput screening,” Synthetic Biology, vol. 8, no. 1, p. ysad012, 2023.

[68] S. Zobel, I. Benedetti, L. Eisenbach, V. de Lorenzo, N. Wierckx, and L. M. Blank, “Tn7-Based Device for Calibrated Heterologous Gene Expression in Pseudomonas putida,” ACS Synthetic Biology, vol. 4, no. 12, pp. 1341–1351, 2015.

[69] A.-M. Anhel, L. Alejaldre, and Á. Goñi-Moreno, “The laboratory automation protocol (lap) format and repository: a platform for enhancing workflow efficiency in synthetic biology,” ACS synthetic biology, vol. 12, no. 12, pp. 3514–3520, 2023.

[70] L. Alejaldre, A. M. Anhel, and Goñi-Moreno, “High-throughput workflow for the genotypic characterization of transposon library variants,” 10.17504/protocols.io.kqdg394jzg25/v1, Oct. 2022.

[71] M. Imakaev, G. Fudenberg, R. P. McCord, N. Naumova, A. Goloborodko, B. R. Lajoie, J. Dekker, and L. A. Mirny, “Iterative correction of Hi-C data reveals hallmarks of chromosome organization,” Nature Methods, vol. 9, pp. 999–1003, Oct. 2012. Publisher: Nature Publishing Group.

[72] J. M. Miró-Bueno and A. Rodríguez-Patón, “A simple negative interaction in the positive transcriptional feedback of a single gene is sufficient to produce reliable oscillations,” PLoS One, vol. 6, no. 11, p. e27414, 2011.

