## Supplemental Info for "Reprogramming genetic circuits using space"

### Supplementary Information
### Reprogramming genetic circuits using space

Lorea Alejaldre[1,2,†], Jesús Miró-Bueno[1,2,†], Angeles Hueso-Gil[1,2], Lewis Grozinger[1,2], Huseyin Tas[1,‡], Sina Geißler[1], and Ángel Goñi-Moreno[1,2,*]

[1]Systems Biology Department, Centro Nacional de Biotecnología (CNB), CSIC, Darwin 3, 28049 Madrid, Spain
[2]Centro de Biotecnología y Genómica de Plantas, Universidad Politécnica de Madrid (UPM)-Instituto Nacional de Investigación y Tecnología Agraria y Alimentaria (INIA/CSIC), 28223, Madrid, Spain
[†]These authors contributed equally to this work.
[‡]Current affiliation: Department of Genetics, Harvard Medical School, 77 Avenue Louis Pasteur, Boston, MA 02115

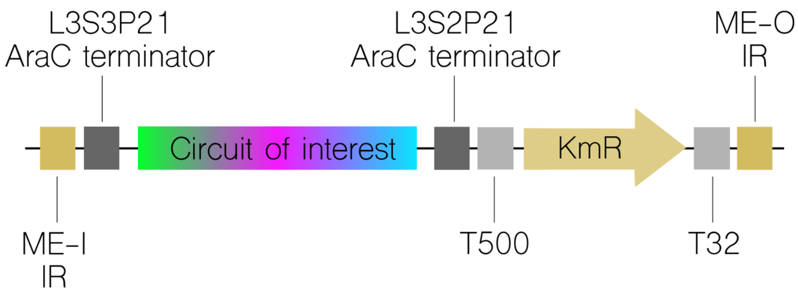


Figure S1: Basic design of insertion cassettes used in this work. Flanked by the insertion regions (ME-I and ME-O when the delivery transposon is Tn5; IR regions when Mariner transposon is used) circuit of interest ($a_m$, $a_0$, $l_m$, $l_0$, $t_m$, $t_0$ or toggle switch) is isolated by the terminators L3S3P21 - L3S2P21 (for $a_m$, $l_m$, $t_m$), L3S3P21 - AraC terminator ($a_0$, $l_0$, $t_0$) or AraC terminator - L3S2P1 (toggle switch), followed by T500, Km resistance selection gene and T32 terminator.

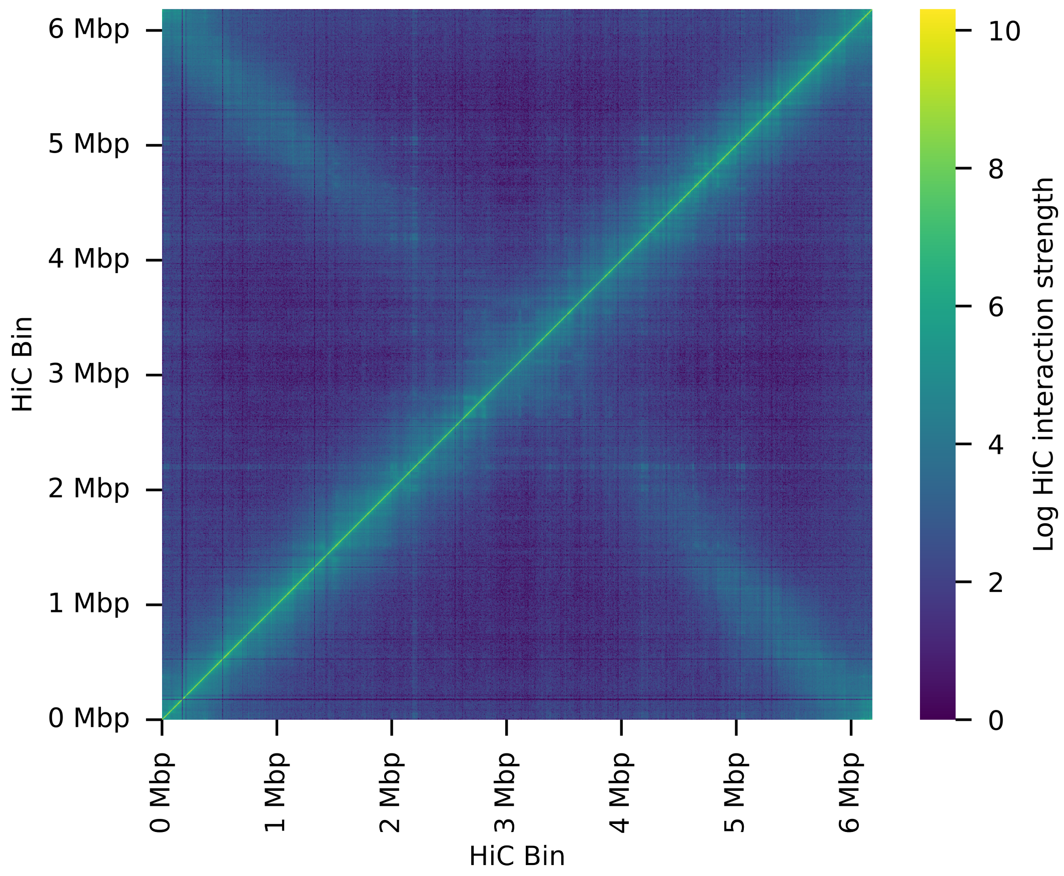


Figure S2: Higher resolution version of Figure 1-C, the HiC contact matrix. Colour indicates the logarithmic interaction strength between two bins, based on number of interactions found in sequencing reads.

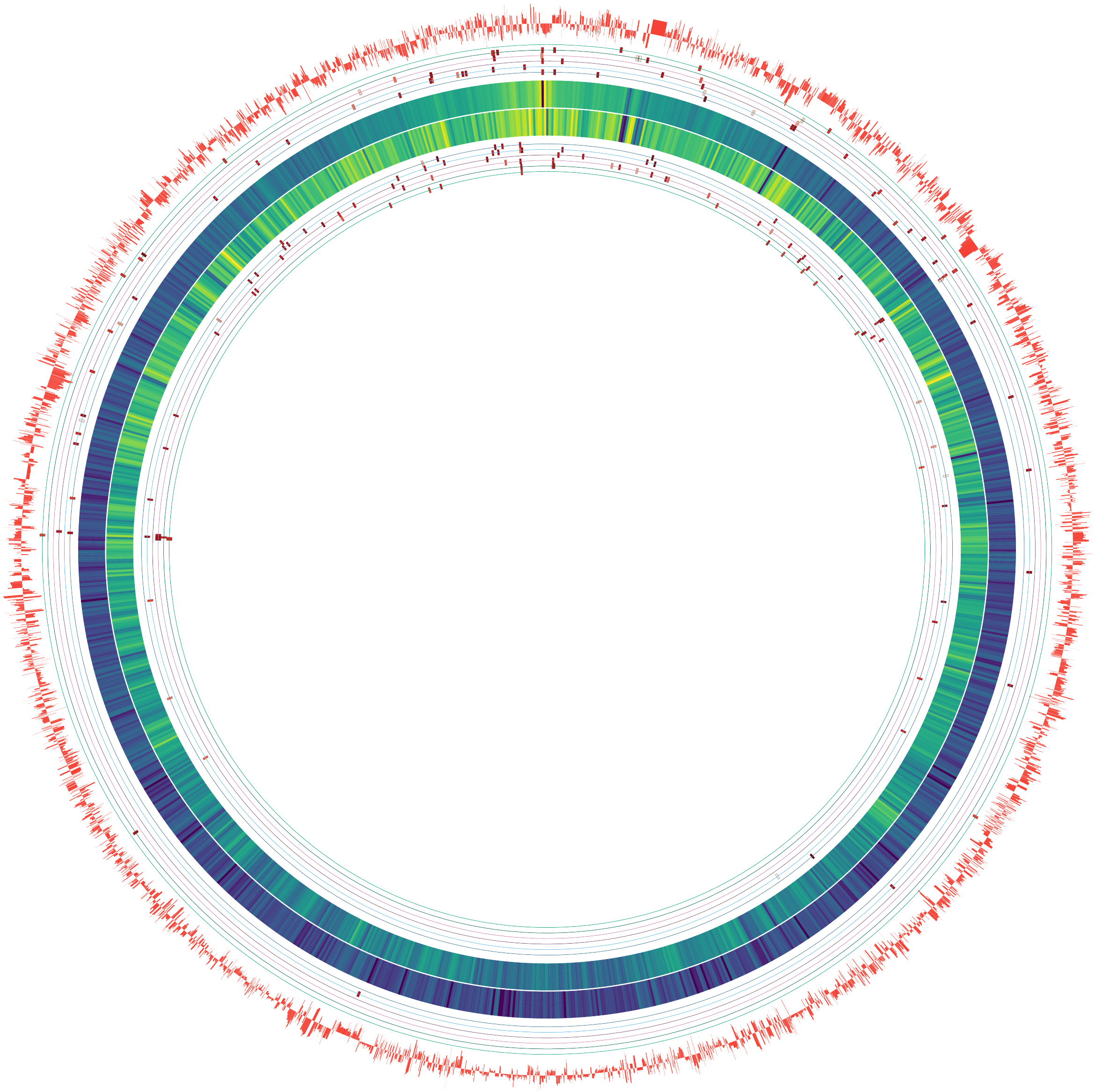

Figure S3: Higher resolution version of Figure 1-D.

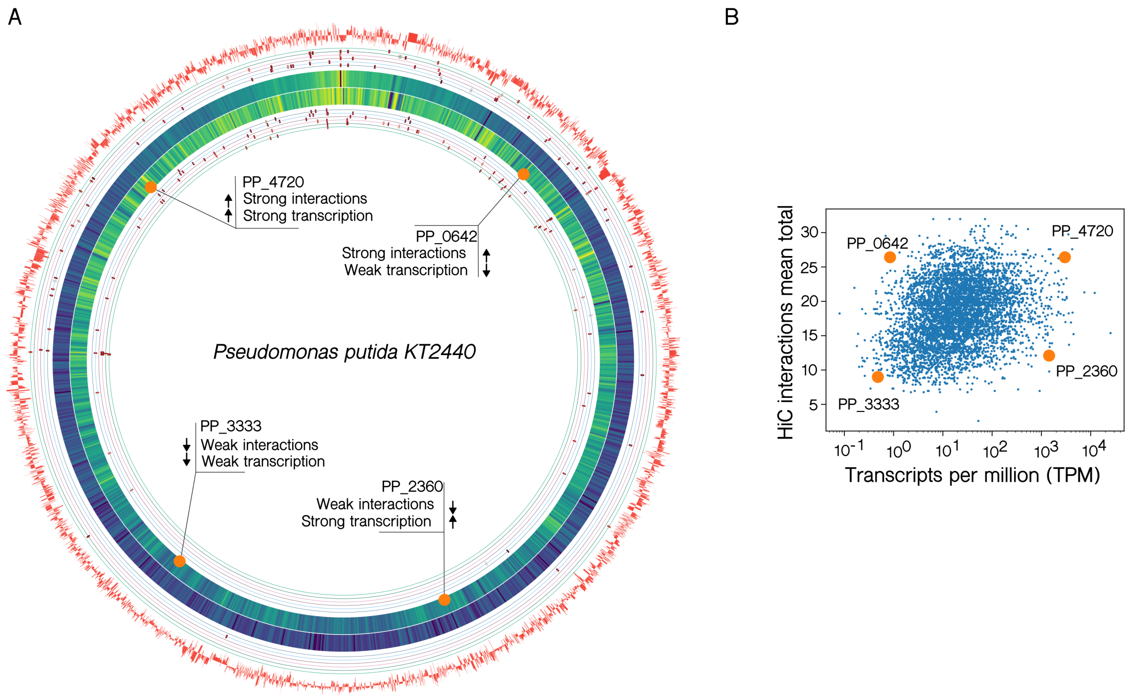


Figure S4: **A)** Four positions are highlighted (orange circles), corresponding to loci with different interaction (HiC) and transcription (RNA-seq) strength **B)** The transcripts per million (TPM) for each loci obtained from RNA-seq analysis is plotted against their mean HiC interaction with all other parts of the chromosome. Orange dots correspond to those highlighted in panel A.

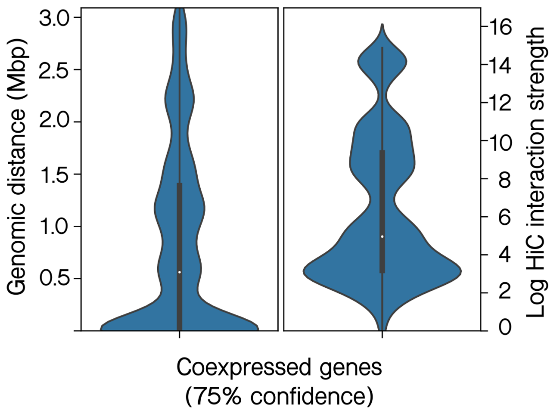


Figure S5: Pairs of coexpressed genes in *P. putida* KT2440, taken from the STRING database. Plots show the distribution of the genomic distance between them in base pairs (left) and the interaction strength between them as taken from the HiC contact matrix. Co-expressed genes are clearly clustered in three large groups.

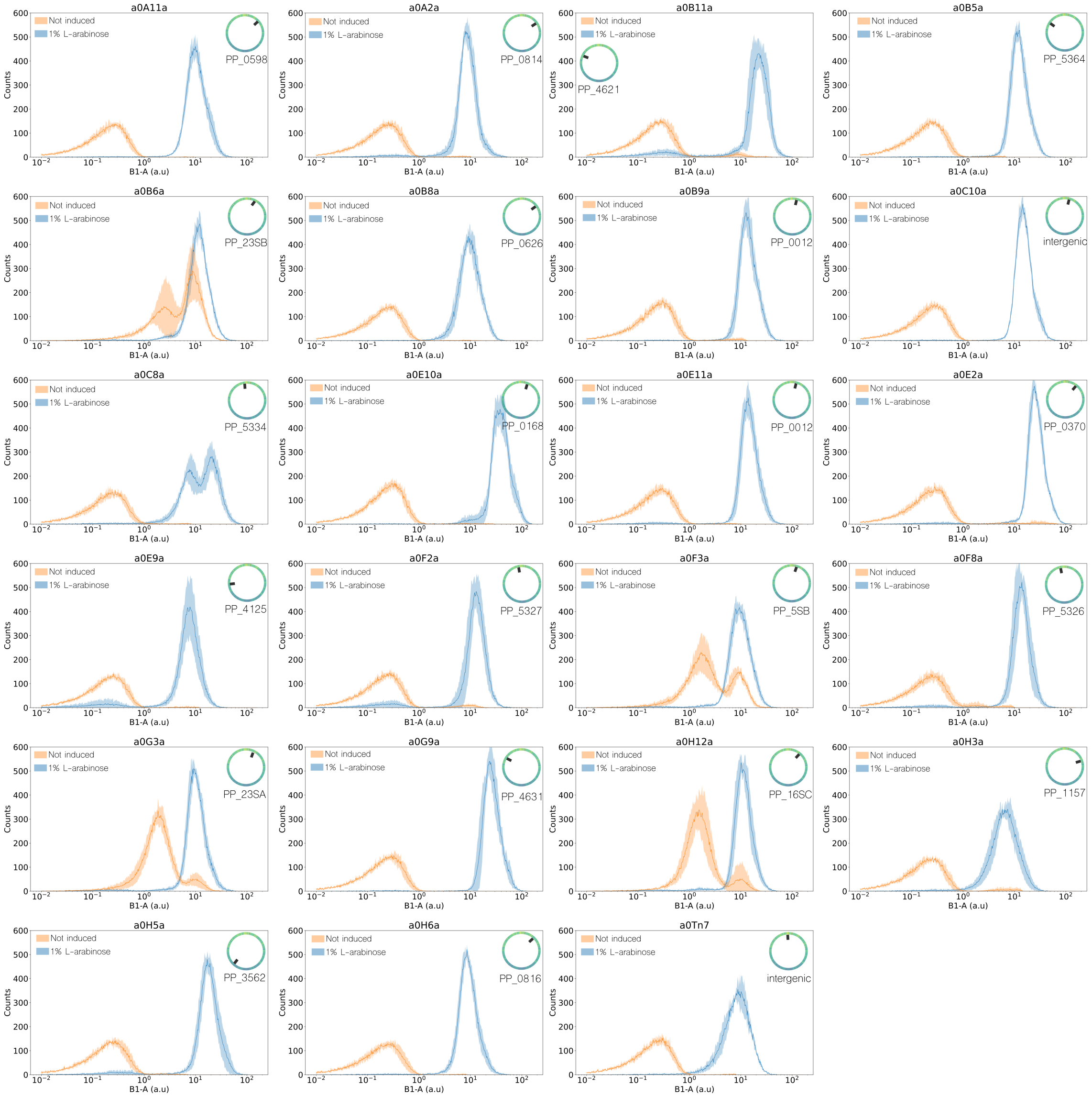


Figure S6: Histograms of $a_0$ library clones in the absence (orange) and presence (blue) of 1% of the inducer L-arabinose after 4 hours of incubation. Position of each insertion is shown next to its fluorescence curves.

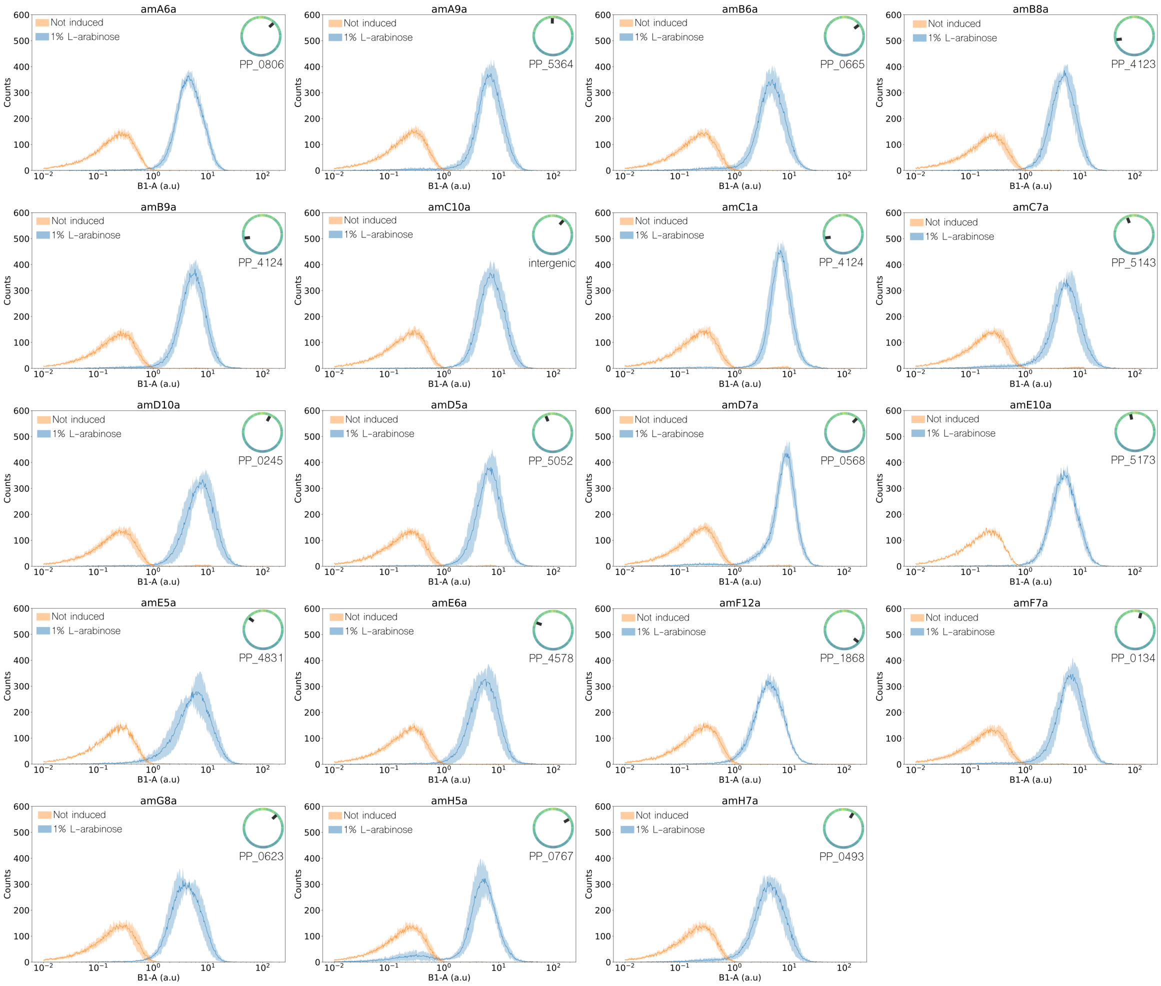


Figure S7: Histograms of a$_m$ library clones in the absence (orange) and presence (blue) of 1% of the inducer L-arabinose after 4 hours of incubation. Position of each insertion is shown next to its fluorescence curves.

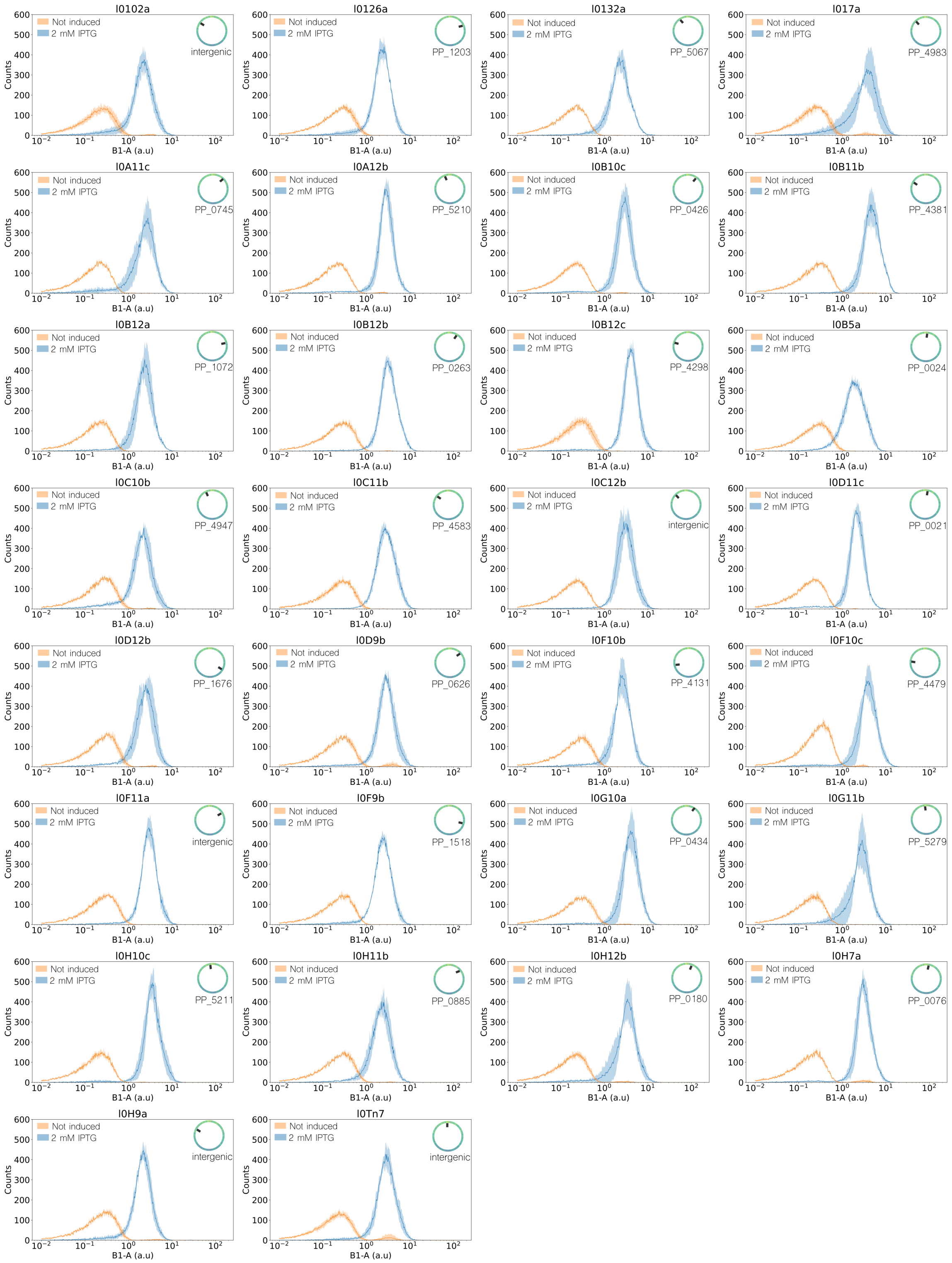


Figure S8: Histograms of $l_0$ library clones in the absence (orange) and presence (blue) of 2 mM of the inducer IPTG after 4 hours of incubation. Position of each insertion is shown next its to fluorescence curves.

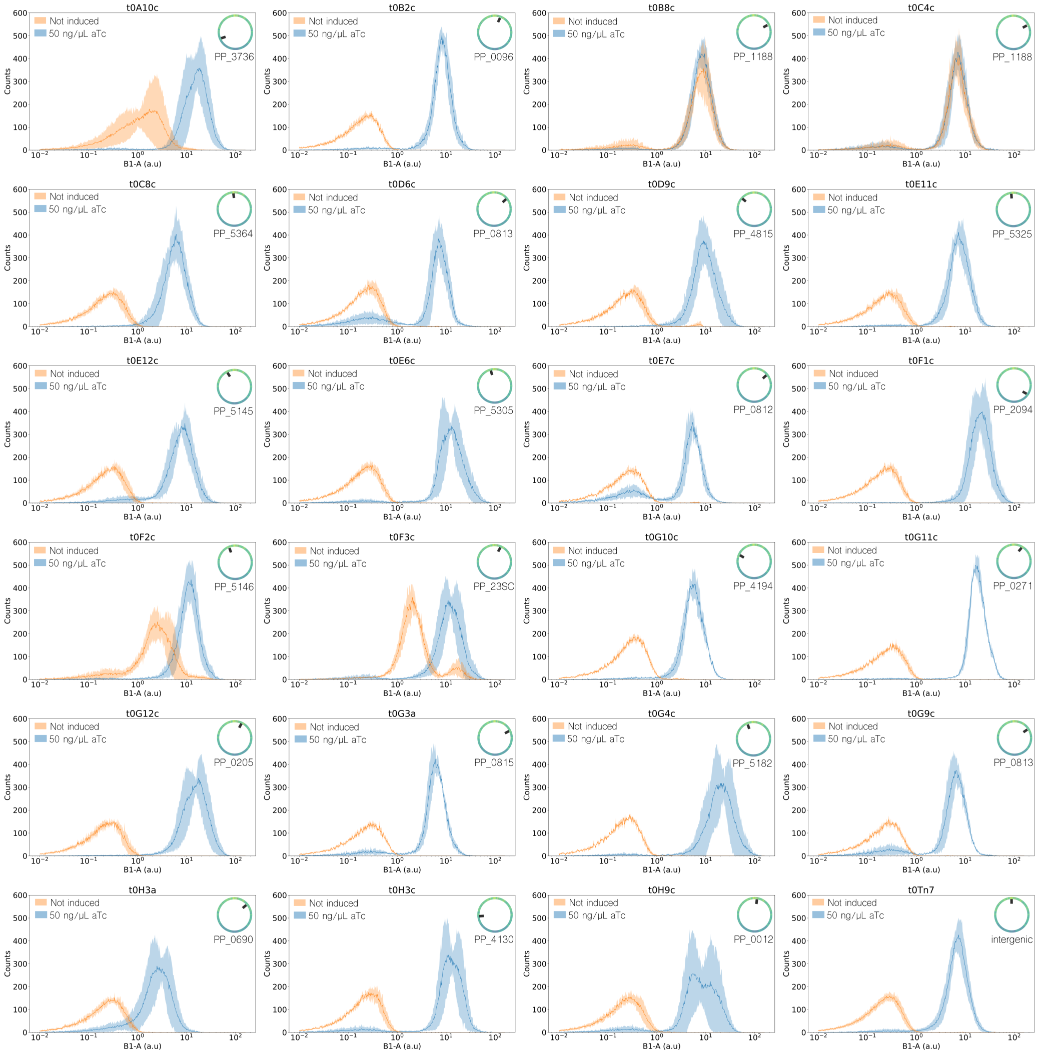


Figure S9: Histograms of $l_m$ library clones in the absence (orange) and presence (blue) of 2 mM of the inducer IPTG after 4 hours of incubation. Position of each insertion is shown next to its fluorescence curves.

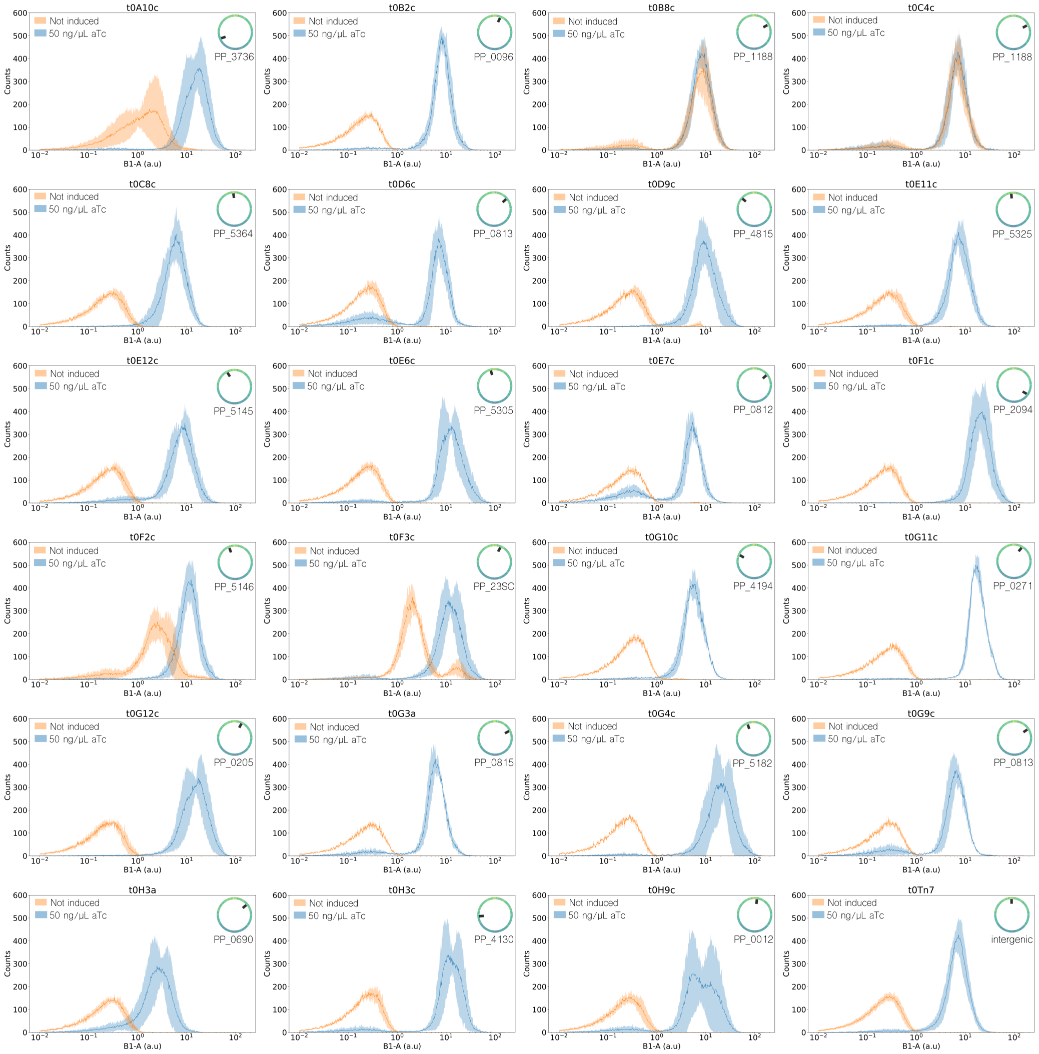


Figure S10: Histograms of $t_0$ library clones in the absence (orange) and presence (blue) of 50 ng/μL of the inducer aTc after 4 hours of incubation. Position of each insertion is shown next to its fluorescence curves.

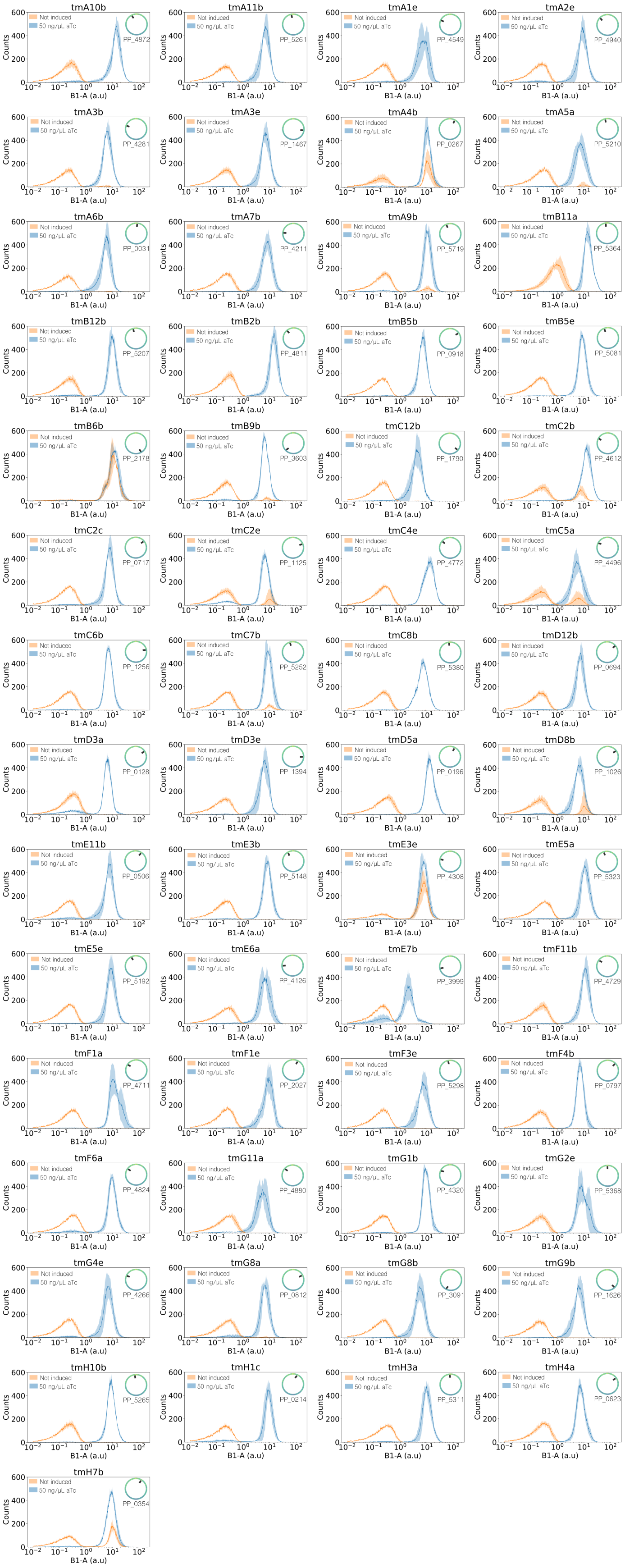


Figure S11: Histograms of $t_m$ library clones in the absence (orange) and presence (blue) of 50 ng/µL of the inducer aTc after 4 hours of incubation. Position of each insertion is shown next to its fluorescence curves.

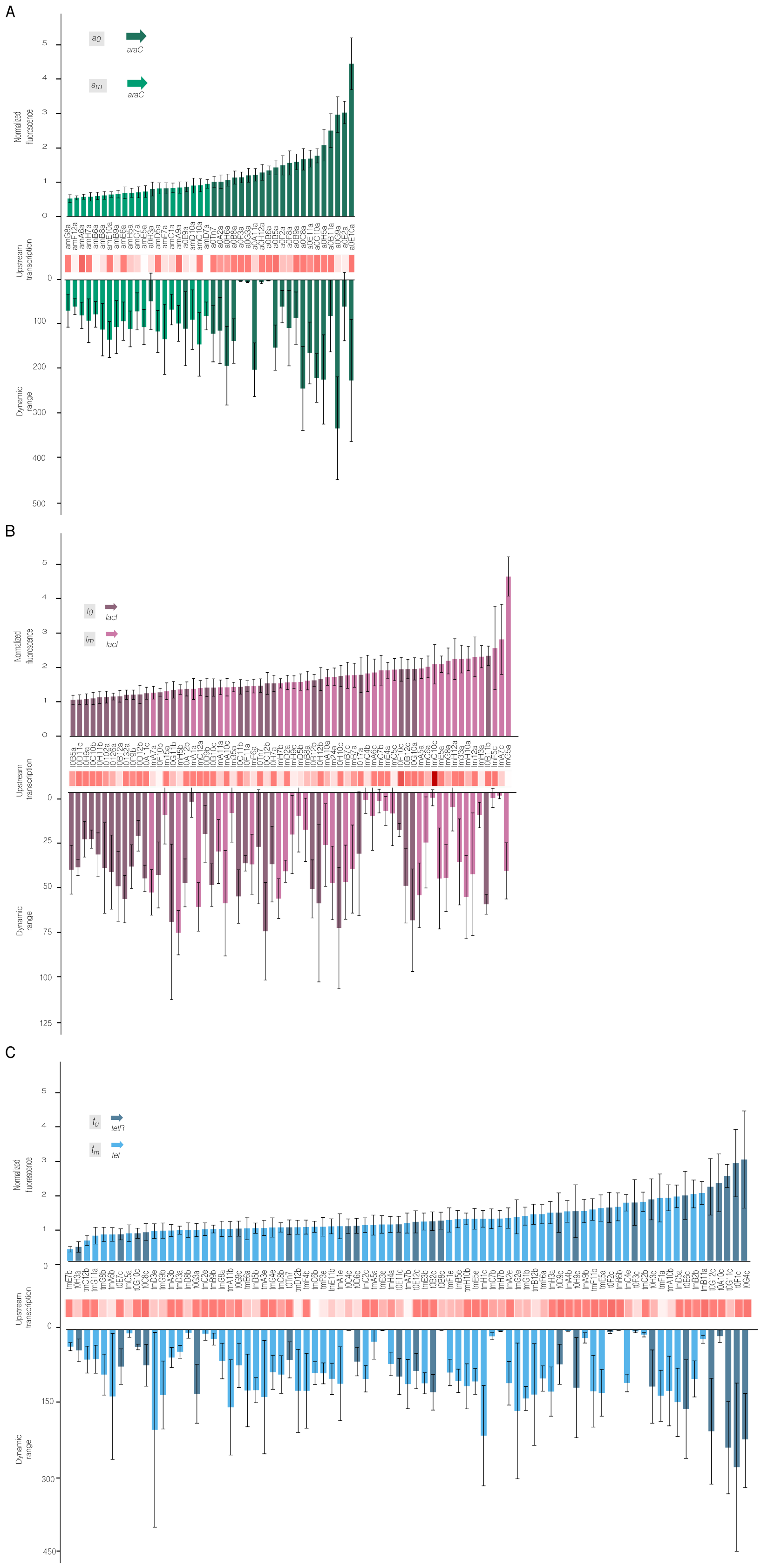


Figure S12: Detailed version of Figure 4 panel C, showing variant names per fluorescent values obtained after 4h incubation with the corresponding inductor, trasnscription level of upstream gene and dynamic range. AraC libraries (A), LacI (B) and TetR (C).

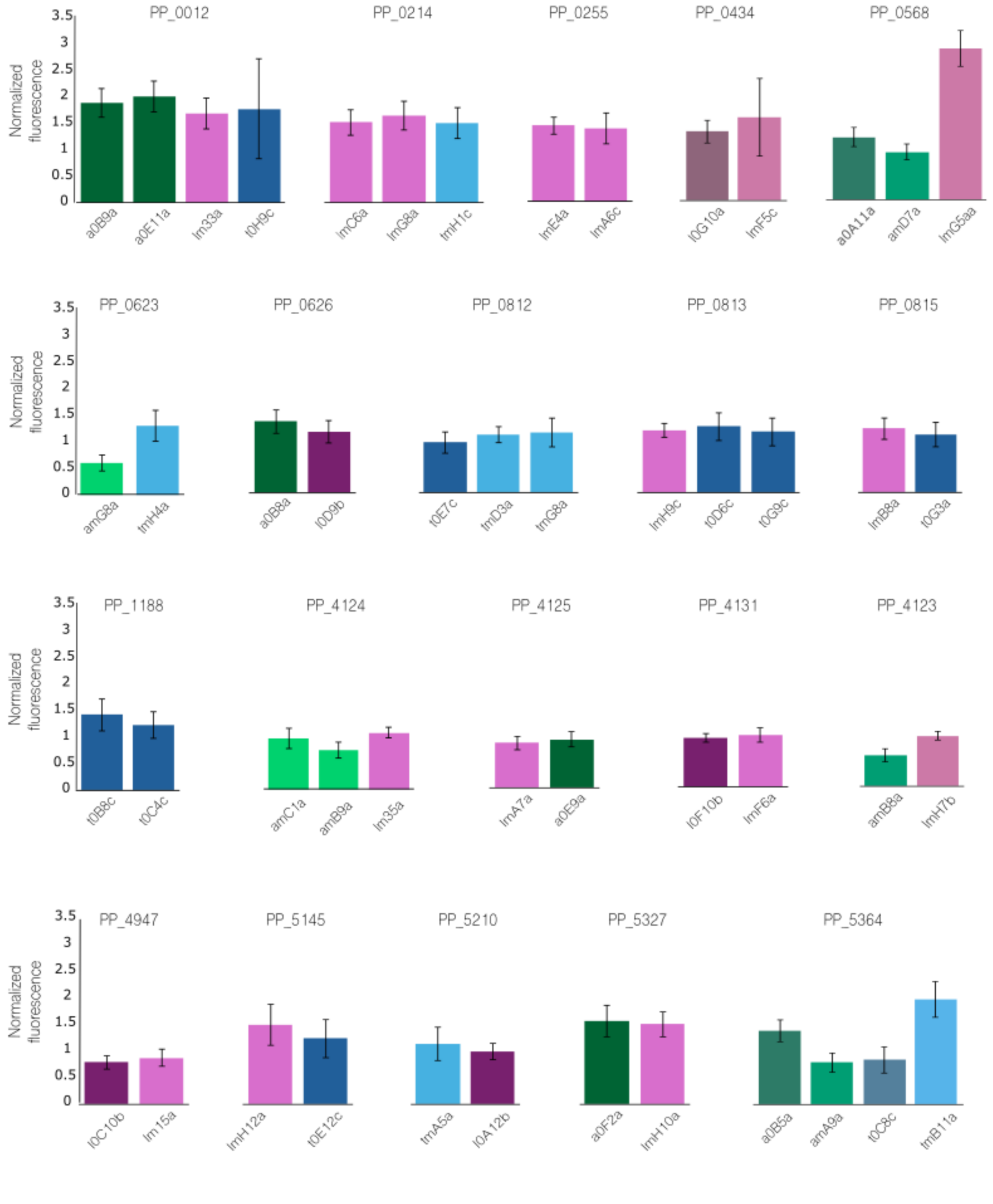


Figure S13: Behaviour in terms of fluorescence produced at saturating induced conditions of the clones that share the same genome position .

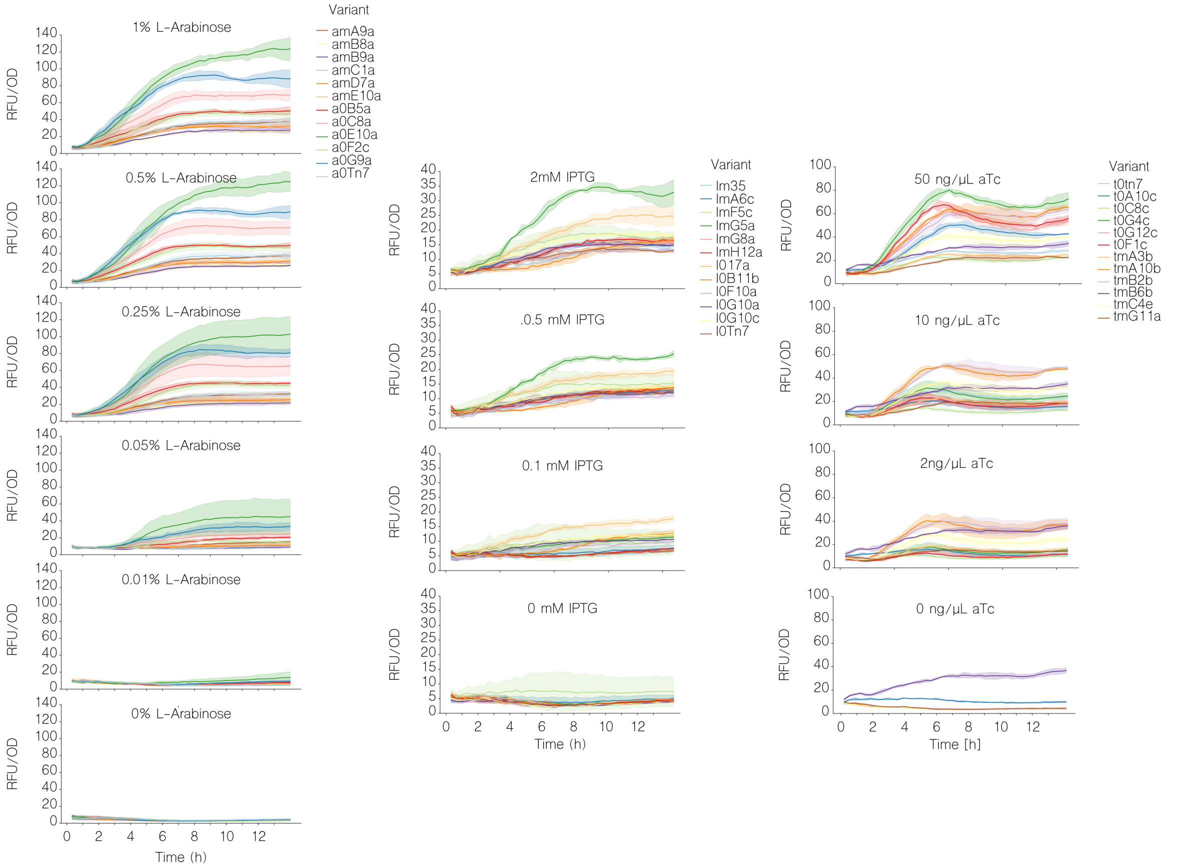


Figure S14: Fluorescence over time normalised by growth for selected variants from AraC (left), LacI (center) and TetR (right) libraries. Each inductor concentration tested is shown in a separate graph for all of the tested variants per library.

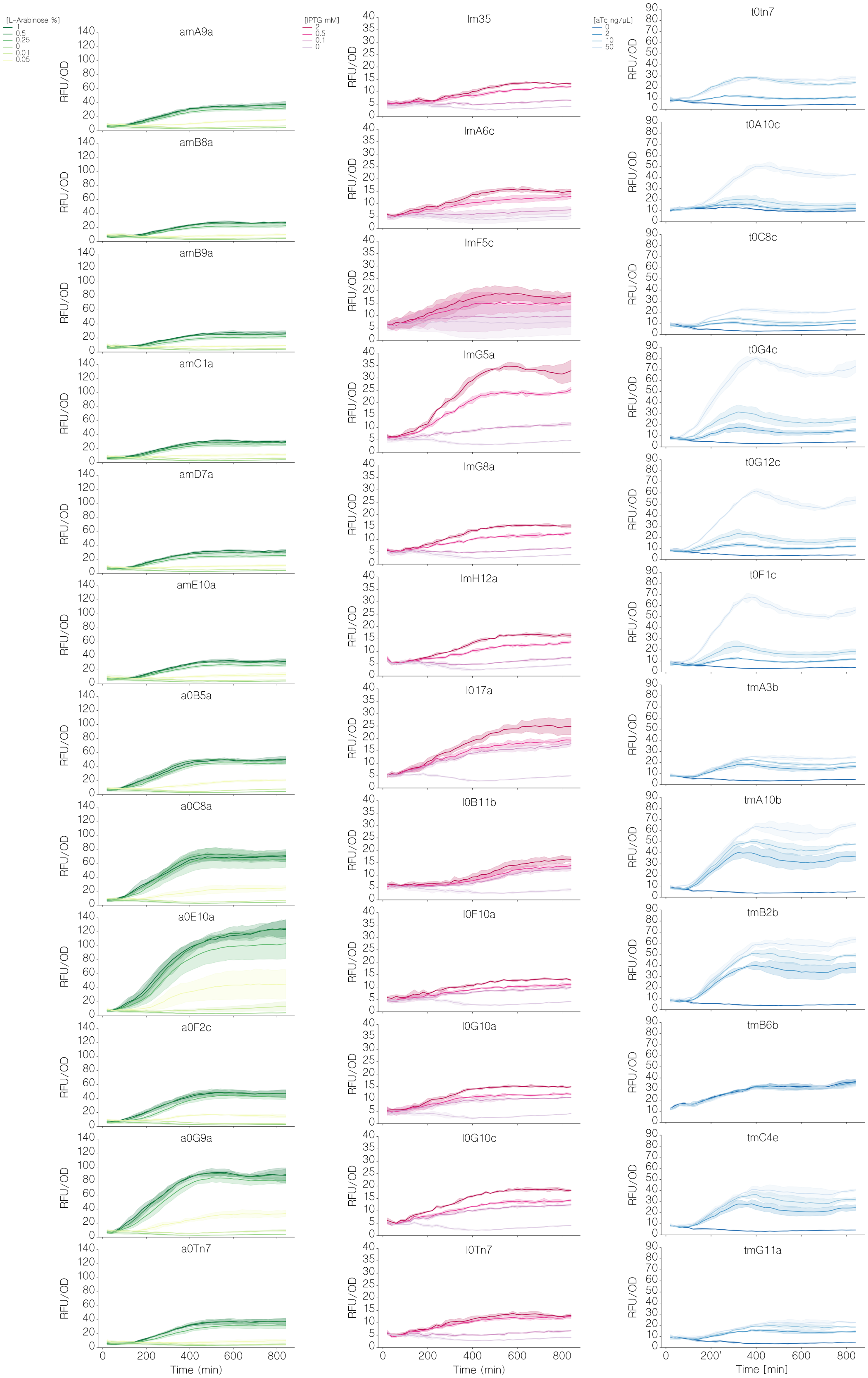


Figure S15: Fluorescence normalised to growth (as optical density) for individual variants of AraC (left - green), LacI (center -magenta) and TetR (right - blue) libraries at different concentrations of inductor.

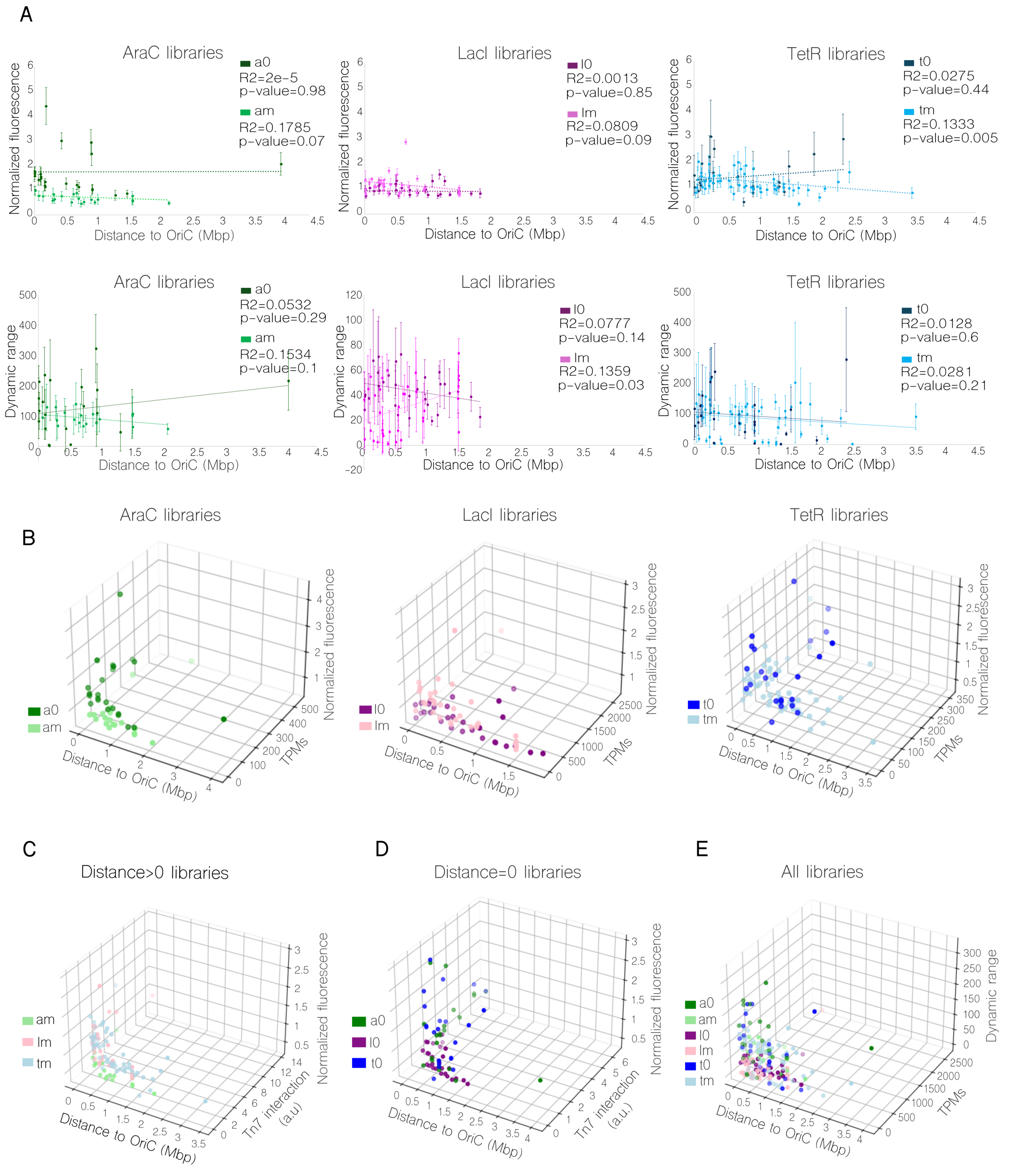


Figure S16: Correlations between selected parameters for the transcriptional cascade libraries. **A)** Correlations for each circuit library (AraC green-left; LacI magenta-center and TetR blue-right) of the normalized fluorescence at inducer saturating conditions and dynamic range of the system with its distance to de OriC. **B)** Similar comparison including the variable transcripts per million (TPM) of the previous gene to the insertion spot of the represented clone. **C)** Normalized fluorescence against distance to OriC and the interaction with *attTn7* position, where the regulator of the circuit is placed, for the libraries $a_m$, $l_m$, $t_m$. Specific clones a0Tn7, l0Tn7 and t0Tn7 not shown in this graph due to its full interaction with Tn7. **D)** Normalised fluorescence with respect to *attTn7* site for $a_0$, $t_0$ and $l_0$ libraries against distance to oriC and interaction with *attTn7* position despite regulator is not placed there in those clones, in order to compare results with those shown on panel C. **E)** Correlation for all the libraries between dynamic range values against distance to oriC and TPMs.

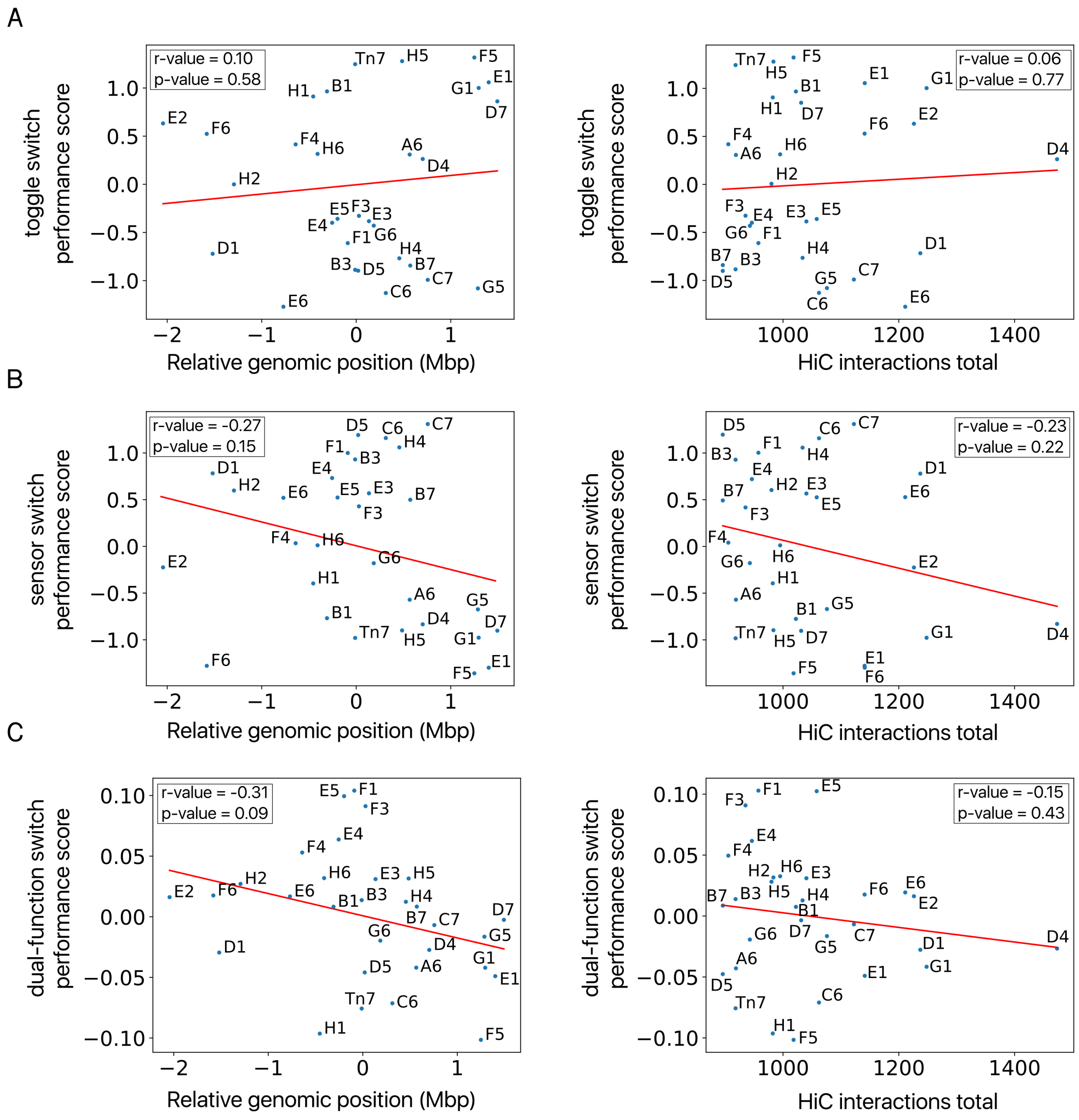


Figure S17: **A)** Toggle, **B)** sensor, and **C)** dual-function switch performance scores plotted against their relative genomic positions (left) and HiC interactions (right). The red line represents the linear regression, with the r-values and p-values displayed in each graph.

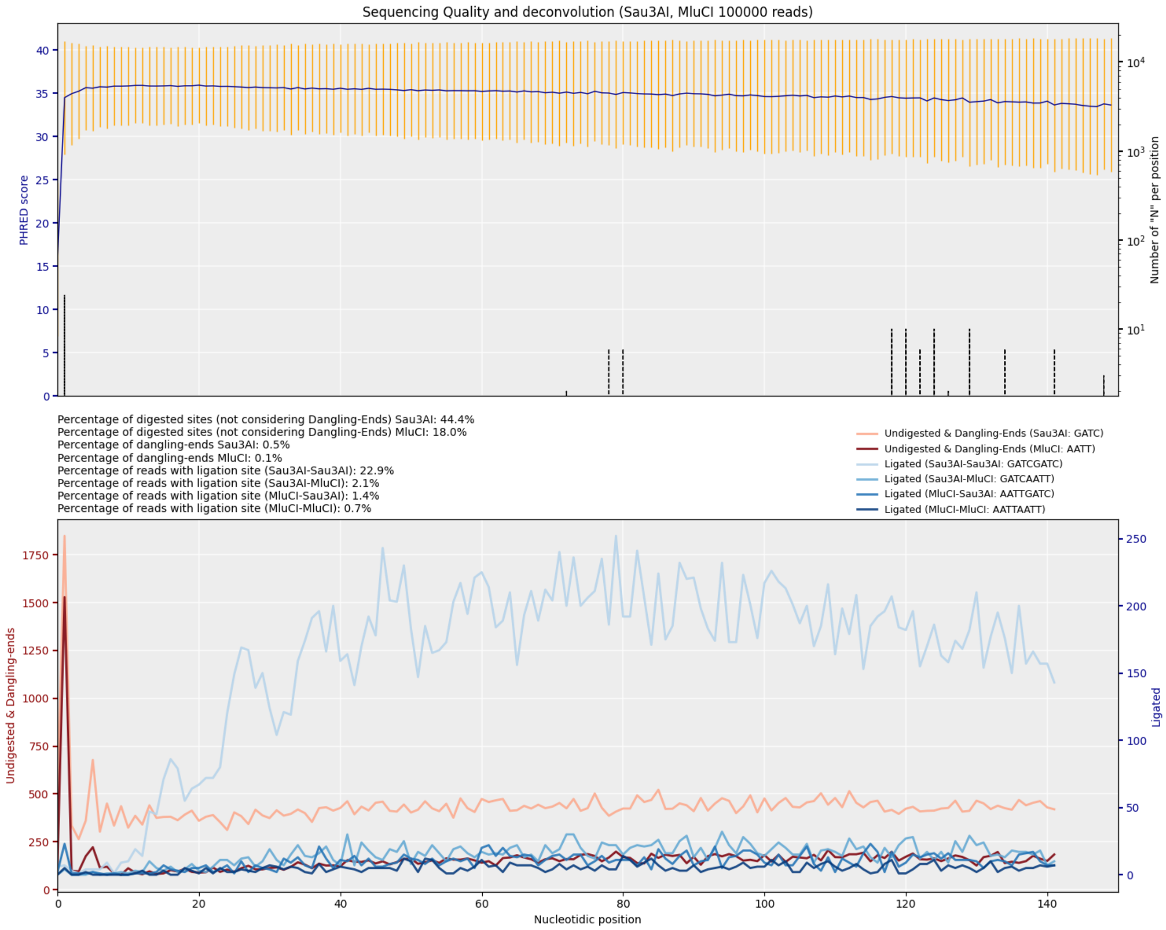


Figure S18: Quality control plot for the first read of the featured wild-type KT2440 sample.

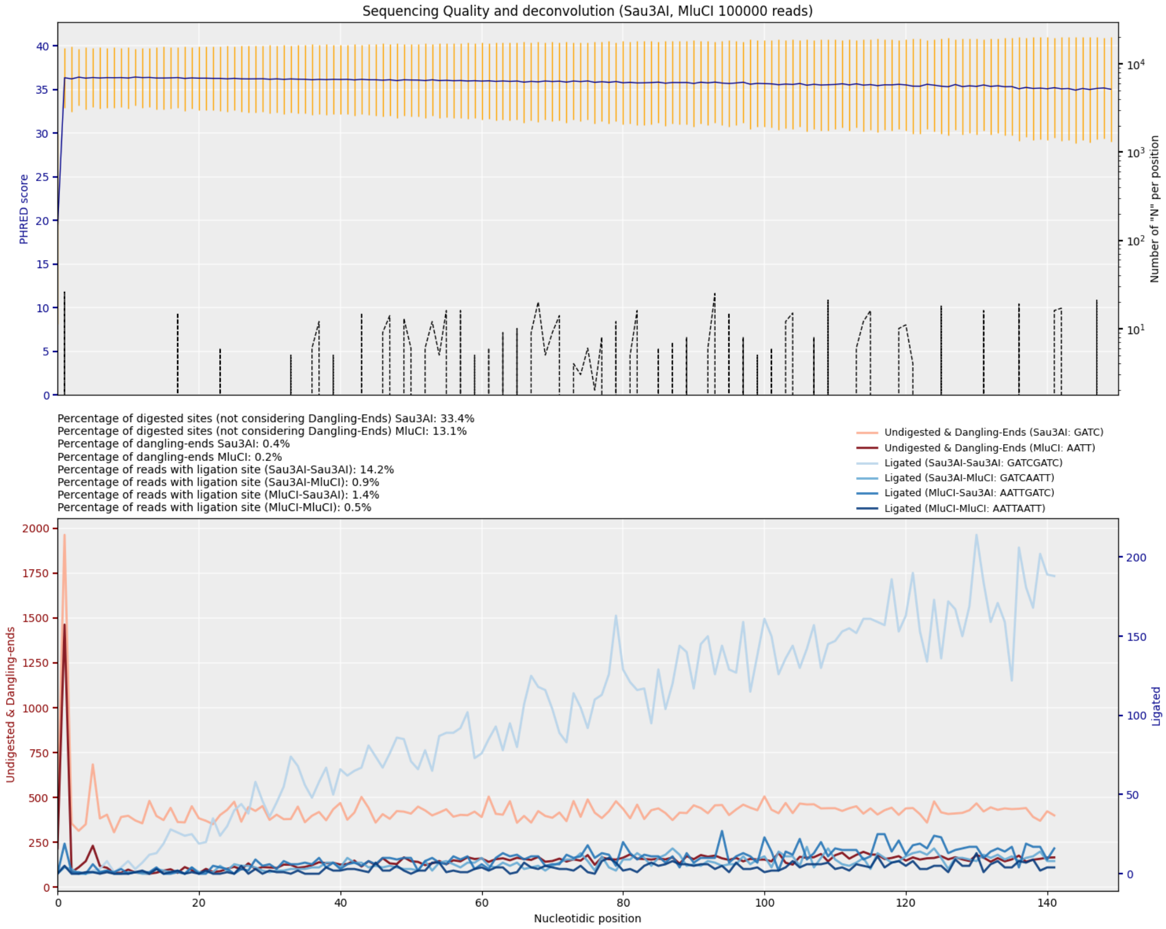


Figure S19: Quality control plot for the second read of the featured wild-type KT2440 sample.

| Clone name | Insertion start | Strand | Feature type | Locus tag |
|---|---|---|---|---|
| TS20D5 | 20744 | plus | CDS | PP_0016 |
| TS20F3 | 28889 | plus | CDS | PP_0024 |
| TS20E3 | 134668 | plus | CDS | PP_0129 |
| TS20G6 | 185462 | plus | CDS | PP_0161 |
| TS20C6 | 312961 | plus | CDS | PP_0257 |
| TS20H4 | 455868 | minus | CDS | PP_0375 |
| TS20H5 | 485061 | minus | intergenic | NA |
| TS20A6 | 565607 | plus | CDS | PP_0481 |
| TS20B7 | 571584 | plus | intergenic | NA |
| TS20D4 | 703407 | minus | CDS | PP_0598 |
| TS20C7 | 756796 | plus | CDS | PP_0646 |
| TS20F5 | 1249629 | minus | CDS | PP_1090 |
| TS20G5 | 1287390 | minus | CDS | PP_1125 |
| TS20G1 | 1295608 | minus | CDS | PP_1132 |
| TS20E1 | 1401762 | minus | CDS | PP_1226 |
| TS20D7 | 1493136 | minus | CDS | PP_1306 |
| TS20E2 | 4137842 | plus | CDS | PP_3641 |
| TS20F6 | 4600389 | plus | CDS | PP_4072 |
| TS20D1 | 4662833 | plus | CDS | PP_4124 |
| TS20H2 | 4888153 | minus | CDS | PP_4297 |
| TS20E6 | 5411477 | minus | CDS | PP_4754 |
| TS20F4 | 5540786 | plus | CDS | PP_4872 |
| TS20H1 | 5726855 | plus | CDS | PP_5025 |
| TS20H6 | 5773060 | plus | intergenic | NA |
| TS20B1 | 5872531 | plus | CDS | PP_5145 |
| TS20E4 | 5926942 | plus | CDS | PP_5195 |
| TS20E5 | 5984165 | minus | CDS | PP_5242 |
| TS20F1 | 6092027 | minus | CDS | PP_5343 |
| TS20B3 | 6170409 | minus | intergenic | NA |
| TS20Tn7 | 6170441 | plus | intergenic | NA |

Table S1: Location of variants for Toggle switch TS20 library

| | **Primer name** | **Sequencence** |
|---|---|---|
| 1 | F1-RF-EcoRI-pTn7 | CGCCTAGGCCGCGGCCGCGCGAATTCAACACCCCTTGTATTACTGT |
| 2 | R1-RF-HindIII-pTn7 | AGTCACGACGCGGCCGCAAGCTTACCTTGCTCATGTTTGACAGC |
| 3 | F1-RF-XmaI-RgFx | GGGCCTTTTTTCGTTTTGGTCCCCCGGGAATGGTCACCAT |
| 4 | R1-RF-HindIII pBAM | GGCTTTGGCGGCCGCAAGCTTACCTTGCTCATGTTTGACAGC |
| 5 | Fwd-EcoRI-RgFx | TGCGACCTAGAATTCAACACCCCTTG |
| 6 | Rev-HindIII-RgFx | CATCACTAAGCTTACCTTGCTCATG |
| 7 | Rev-XmaI-RgFx | ATGGTGACCATTCCCGGG |
| 8 | Fwd-XmaI-RgFx | TTTGGTCCCCCGGGAATG |
| 9 | Fwd-EcoRI-Toggle | TCGACCTAGAATTCTGCTCATGTTTG |
| 10 | Rev-HindIII-Toggle | TAGACTAAGCTTGGACCAAAACG |
| 11 | R1-RF-HindIII-pBAMD1-2 | GGCTTTGGCGGCCGCAAGCTTACCTTGCTCATGTTTGACAGC |
| 12 | F1-RF-XmaI | GCGCGAATTCGAGCTCGGTACCCGGGAATGGTCACCAT |
| 13 | F1-QC-MmeI-TetR | GAGGTCGG**C**ATCGAAGGTTTAACAACCCGTAAACTC |
| 14 | R1-QC-MmeI-TetR | CTTCGAT**G**CCGACCTCATTAAGCAGCTCTAATGCG |

Table S2: List of primers used in this study

| Plasmid | Strategy | Primers | Template | Backbone | Restriction Enzymes |
|---|---|---|---|---|---|
| pTn7-LacI | Traditional cloning | Fwd-XmaI-RgFx/Rev-HindIII-RgFx | IPTG-LacI-pTac | pTn7-M | XmaI/HindIII |
| pTn7-IPTG-LacI-pTac-YFP | Traditional cloning | Fwd-EcoRI-RgFx/Rev-HindIII-RgFx | IPTG-LacI-pTac | pTn7-M | EcoRI/HindIII |
| pTn7-TetR | Gibson Assembly | F1-RF-XmaI-RgFx/R1-RF-HindIII pBAM | aTc-TetR-pTet | pTn7 | XmaI/HindIII |
| pTn7-aTc-TetR-pTet-YFP | Gibson Assembly | F1-RF-EcoRI-pTn7/R1-RF-HindIII-pTn7 | aTc-TetR-pTet | pTn7 | EcoRI/HindIII |
| pTn7-AraC | Gibson Assembly | F1-RF-XmaI/R1-RF-HindIII-pTn7 | Lara-AraC-pBAD | pTn7-M | XmaI/HindIII |
| pTn7-Lara-AraC-pBAD-YFP | Gibson Assembly | F1-RF-EcoRI-pTn7/R1-RF-HindIII-pTn7 | Lara-AraC-pBAD | pTn7 | EcoRI/HindIII |
| pTn7-TS2-msfGFP-mKate2 | Traditional cloning | Fwd-EcoRI-Toggle/Rev-HindIII-Toggle | SOE PCR from TS2-LacI-mKate2 and TS2-TetR-msfGFP | pTn7-M | EcoRI/HindIII |
| pBLAM1.2-TS2-msfGFP-mKate2 | Traditional cloning | Fwd-EcoRI-Toggle/Rev-HindIII-Toggle | SOE PCR from TS2-LacI-mKate2 and TS2-TetR-msfGFP | pTn7-M | EcoRI/HindIII |
| pBLAM1.2-TS2MmeI | Whole-plasmid site-directed mutagenesis | F1-QC-MmeI-TetR/R1-QC-MmeI-TetR | pBLAM1.2-TS2-msfGFP-mKate2 | pBLAM1-2-TS2-msfGFP-mKate2 | DpnI |
| pBAMD1.2-pTac-YFP | Traditional cloning | Fwd-EcoRI-RgFx/Rev-XmaI-RgFx | IPTG-LacI-pTac | pBAMD1.2 | EcoRI/XmaI |
| pBAMD1.2-IPTG-LacI-pTac-YFP | Traditional cloning | Fwd-EcoRI-RgFx/Rev-HindIII-RgFx | IPTG-LacI-pTac | pBAMD1-2 | EcoRI/HindIII |
| pBLAM1.2-IPTG-LacI-pTac-YFP | Traditional cloning | Fwd-EcoRI-RgFx/Rev-HindIII-RgFx | IPTG-LacI-pTac | pBLAM1-2 | EcoRI/HindIII |
| pBAMD1.2-pTet-YFP | Traditional cloning | Fwd-EcoRI-RgFx/Rev-XmaI-RgFx | aTc-TetR-pTet | pBAMD1-2 | EcoRI/XmaI |
| pBAMD1.2-aTc-TetR-pTet-YFP | Gibson Assembly | F1-RF-XmaI-RgFx/R1-RF-HindIII-pTn7 | aTc-TetR-pTet | pBAMD1.2-pTet-YFP | XmaI/HindIII |
| pBLAM1.2-aTc-TetR-pTet-YFP | Gibson Assembly | F1-RF-EcoRI/R1-RF-HindIII-pBAMD1-2 | aTc-TetR-pTet | pBLAM1-2 | EcoRI/HindIII |
| pBAMD1.2-pBAD-YFP | Traditional cloning | Fwd-EcoRI-RgFx/Rev-XmaI-RgFx | Lara-AraC-pBAD | pBAMD1-2 | EcoRI/XmaI |
| pBAMD1.2-Lara-AraC-pBAD-YFP | Gibson Assembly | F1-RF-EcoRI/R1-RF-HindIII-pBAMD1-2 | Lara-AraC-pBAD | pBAMD1-2 | EcoRI/HindIII |
| pBLAM1.2-Lara-AraC-pBAD-YFP | Gibson Assembly | F1-RF-EcoRI/R1-RF-HindIII-pBAMD1-2 | Lara-AraC-pBAD | pBLAM1-2 | EcoRI/HindIII |

Table S3: Cloning method description for plasmids constructed in this study
